# Widespread *Saccharomyces* hybridization in agave fermentations arises repeatedly from asymmetric parental diversity

**DOI:** 10.64898/2026.11.04.764348

**Authors:** J. Abraham Avelar-Rivas, Xitlali Aguirre-Dugua, Claudio López-Gallegos, Iván Sedeño, Ricardo Echavarría-Solana, Erick Santiago-Garduño, Mariela Arias, Manuel R. Kirchmayr, Gianni Liti, Alexander DeLuna, Lucia Morales, Eugenio Mancera

## Abstract

Interspecific hybridization can profoundly reshape genomes by bringing together divergent genetic backgrounds, yet its ecological and evolutionary dynamics remain poorly understood at the population level. Here, we identify an extensive hybridization system in traditional agave fermentations across Mexico involving *Saccharomyces cerevisiae*, a species frequently associated with human-managed environments, and its predominantly wild relative *S. paradoxus*. A genomic survey reveals unexpectedly high hybrid frequencies in geographically distant regions with contrasting environments. Targeted sampling of distilleries and surrounding natural sites reveals a strong ecological bias, with hybrids recovered almost exclusively from distilleries, although both parental species occur in anthropogenic and natural habitats. Hybrid genomes show extensive architectural diversity, including distinct ploidies, variable parental genome contributions, variable patterns of heterozygosity, and contrasting mitochondrial ancestries. Phylogenomic and population-structure analyses of hybrid subgenomes demonstrate multiple independent origins. These recurrent events show pronounced asymmetry in parental genetic diversity. Hybrids derive from several *S. cerevisiae* populations, already shaped by extensive introgression, whereas every *S. paradoxus* subgenome traces to a single genetically differentiated *S. paradoxus* lineage found exclusively in distilleries, an unexpected association for a predominantly wild species. These findings establish agave fermentations as an exceptional system for studying recurrent interspecific hybridization and reveal how pronounced asymmetry in parental ancestry can shape hybrid formation.

**Impact statement:** Interspecific hybridization is a powerful source of evolutionary novelty, but its dynamics are rarely resolved at the population level outside experimental systems. We uncover a widespread system of recurrent *Saccharomyces* hybridization associated with agave fermentations, revealing multiple independent origins and a striking asymmetry in the parental lineages that contribute to hybrid formation, with diverse *S. cerevisiae* backgrounds repeatedly mating with a single, human-associated *S. paradoxus* lineage.

## Introduction

Hybridization between species can generate genetic combinations that are inaccessible within either parental species, providing an important source of evolutionary novelty. By bringing together divergent genomes, hybridization can introduce adaptive alleles across species boundaries, generate transgressive phenotypes, and create new combinations of traits on which selection can act^1–5^. When hybrids backcross with one of the parental lineages, genetic material from the other parent can persist long after the original hybridization event through introgressions^6–10^. The evolutionary consequences of interspecies hybridization are evident across the tree of life, where gene flow between related species has contributed to adaptation in numerous lineages^5,11–15^. Humans provide a prominent example, whereby admixture with Neanderthals and Denisovans resulted in introgressed genomic regions that influenced adaptation as well as disease susceptibility in modern human populations^16–18^. Hybridization can therefore generate outcomes ranging from transient hybrid genomes to persistent introgression and the emergence of new evolutionary lineages.

Despite its evolutionary importance, the fate of interspecific hybrids remains difficult to resolve in natural populations. Hybridization was long regarded as an exception to species evolution, with interspecific hybrids often considered rare, poorly fertile, and evolutionary dead ends^19,20^. Although this view has changed substantially, most natural systems still lack genomic sampling dense enough to connect hybrids with the diversity of their parental populations. As a result, it remains difficult to determine how persistent contact among genetically diverse parental and hybrid populations gives rise to distinct evolutionary outcomes, including recurrent hybrid formation, diversification of hybrid lineages and introgression into parental populations. Systems in which multiple contemporary hybrids can be studied alongside extant parental populations are therefore especially valuable for understanding hybrid evolution.

The genus *Saccharomyces* provides a powerful system for population-level studies of interspecific hybridization. Thousands of sequenced genomes capture extensive diversity across natural and human-associated environments, while genetic tractability allows genomic observations to be tested experimentally^8,9,21–25^. Foundational population studies established a broad ecological contrast between *S. cerevisiae*, increasingly recognized as a species strongly associated with anthropogenic environments, and its closest relative *S. paradoxus*, regarded primarily as a wild species structured among natural populations^26,27^. Species within the genus readily form interspecific hybrids, and hybridization has contributed to their evolution at different timescales^8,9,10,21,24,27^. An ancient hybridization event underlies the whole-genome duplication shared by *Saccharomyces* and related post-WGD lineages^29^, whereas recent hybrids are especially prominent in human-associated fermentations^30^. The best-known example is the *S. cerevisiae x S. eubayanus* lager yeast *S. pastorianus*^31–33^.

*Saccharomyces* has also revealed how hybrid genomes can evolve after formation, including how loss of heterozygosity (LOH) can restore meiotic recombination and fertility, providing a route to subsequent introgression^9^. Importantly, genomic fragments derived from *S. paradoxus* occur across many *S. cerevisiae* populations and can constitute substantial fractions of the genome in highly introgressed lineages, including Neotropical and Alpechin clades^10,21,22,25^. Yet, complete *S. cerevisiae* x *S. paradoxus* hybrids retaining substantial fractions of both parental genomes remain remarkably rare, with only a handful of environmental isolates described^8,9,25^. This contrast between extensive historical gene flow and the rarity of extant hybrids makes contemporary hybrid populations especially informative for reconstructing how interspecific gene flow originates and evolves.

Here, we reveal the population-level dynamics of *S. cerevisiae* x *S. paradoxus* hybridization associated with open agave fermentations across Mexico. Genomic surveys and targeted sampling of distilleries and surrounding natural environments revealed hybrids at unexpectedly high frequencies, along with a strong association with distilleries and extensive variation in hybrid overall genome architecture. Population-genomic analyses of their parental subgenomes showed that these hybrids arose through multiple independent events and revealed a striking asymmetry in parental contributions: while several *S. cerevisiae* populations contributed to hybrid formation, all *S. paradoxus* ancestry traced to a single, genetically differentiated, human-associated lineage—an unexpected ecological association for this predominantly wild species. These findings establish agave fermentations as a population-level system for investigating recurrent hybridization and hybrid genome evolution against a background already bearing introgressions, a signature of past hybridization.

## Results

### 1. Interspecies hybrids are prevalent and differentially distributed across a broad geographical region

Given the large number of introgressions found in the *S. cerevisiae* strains from Mexico^10,21^, we searched for hybrid isolates by aligning the genomic reads to a concatenated sequence of the reference genomes of all known *Saccharomyces* species. We analyzed a total of 191 *Saccharomyces* genomes from strains previously sampled in agave fermentation tanks^34^ including 42 newly sequenced isolates (**Supplementary Data 1** and **Supplementary Data 2**). Strains in which more than 12.5% (the expected percentage following two backcrosses of an F1 hybrid to one of the parental lineages) of the reads aligned to a non-*S. cerevisiae* genome were classified as hybrids. From the 191 *Saccharomyces* genomes that we analyzed from 113 fermentation tanks across 63 distilleries (**Supplementary Data 3**), 17.2% (n = 33) were hybrids, all between *S. cerevisiae* and *S. paradoxus*. Among the non-hybrid strains, 149 aligned predominantly to the *S. cerevisiae* reference genome and 9 to *S. paradoxus*. None of the strains showed significant read alignment to the genomes of other *Saccharomyces* species.

Hybrid distribution was highly uneven across Mexico; in two of the producing regions of the center and west of Mexico (Central Highlands and West I) the hybrids represented close to 40% of the total analyzed strains (**Fig. 1A**). These strains were isolated from distilleries separated by up to 2,033 km with altitudes ranging from 385 to 2,551 m a.s.l. across agave-producing regions. In this initial genomic survey, we did not identify any hybrid strains in the southernmost region (South Central), West II, or in the Northeast, even though the *S. cerevisiae* isolates from the latter region contain the highest number of introgressions^10^. Hybrids were also recovered in the Northwest (3/16; 18.8%) and in the Balsas Basin (9/36; 25%). These regional differences were significant among isolates with complete location metadata (*n* = 191; two-sided Fisher’s exact test, *p* < 10^-5^).

**Figure 1.**
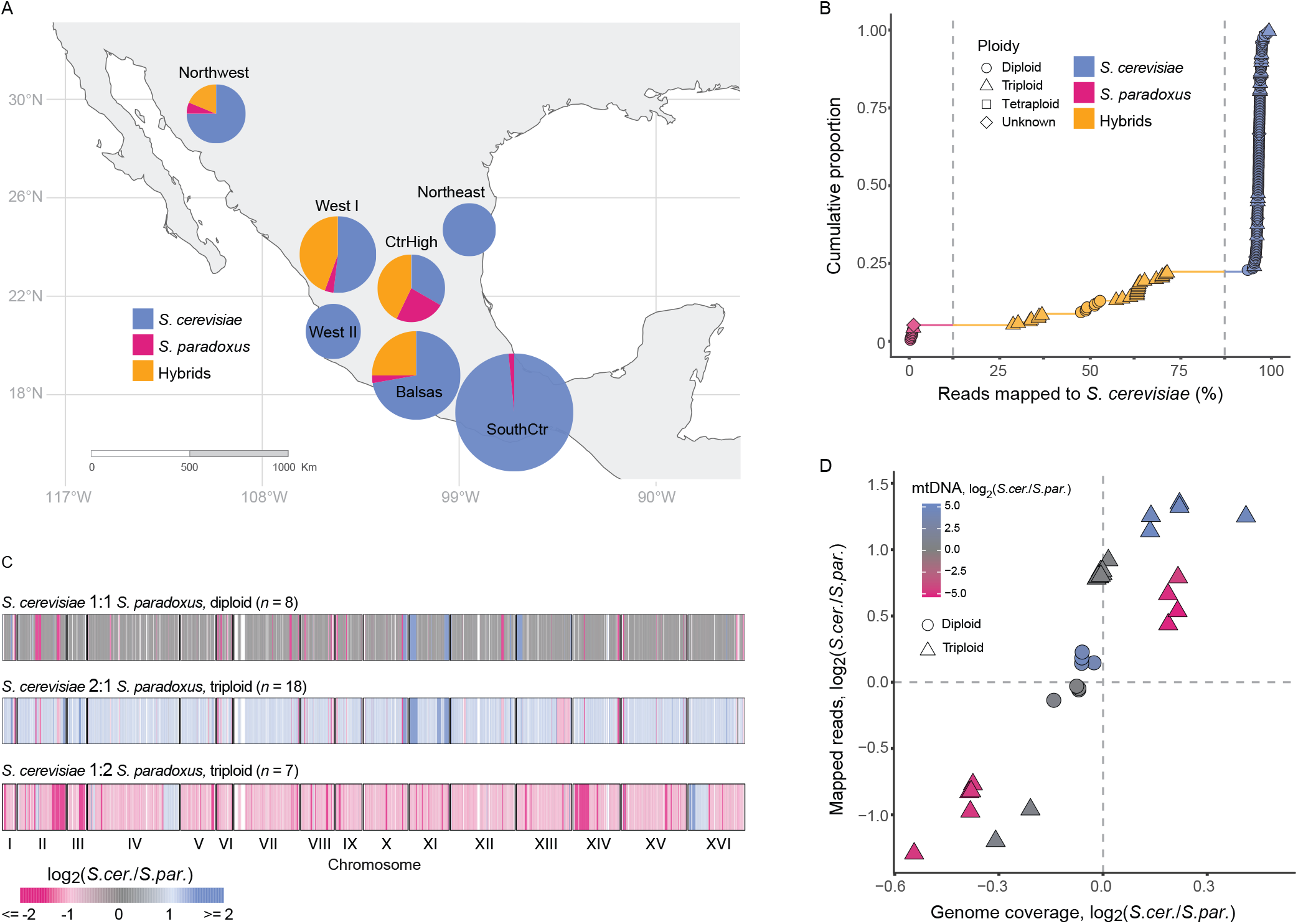
*S. cerevisiae* x *S. paradoxus* hybrids are common in agave fermentations from specific regions of Mexico and exhibit substantial genomic diversity. A) Frequencies of hybrid and parental-species strains across the seven agave-spirit-producing regions of Mexico (Central Highlands [CtrHigh]; South Central [SouthCtr]). Only strains from the first Results section are included. B) Cumulative distribution of the genomic composition of the analyzed strains, shown as the percentage of reads mapping to each parental species genome. Strains were classified as hybrids (yellow) if more than 12.5% of their reads mapped to the genome of the alternate parental species. Ploidy of the strains is depicted by the shape of the dots as specified in the legend. C) Subgenome composition along chromosomes of three hybrid strains representative of the most common genomic configurations. Vertical segments represent genetic markers colored according to their 1-kb depth coverage ratio as depicted in the legend. D) Scatter plot illustrating the genomic diversity of hybrid strains based on parental nuclear genome coverage (log_2_ *S. cerevisiae*/*S. paradoxus*), parental nuclear genome content measured as the number of reads mapping to each parental genome (log_2_ *S. cerevisiae/S. paradoxus*), ploidy (indicated by symbol shape), and mitochondrial composition measured as the number of reads mapping to each parental mitochondrial genome (log₂ *S. cerevisiae/S. paradoxus*) and shown by the symbol color.

The fraction of the reads that aligned to the *S. paradoxus* genome in the hybrid isolates varied considerably, from 28.1% to 70.5% (**Fig. 1B**). To characterize this variation further, we experimentally and computationally determined the ploidy of the hybrids (**Fig. 1B, C, D**). Diploid hybrids with a 1:1 *S. cerevisiae:S. paradoxus* subgenome ratio along most of the genome were a minority (8/33; 24.2% of the hybrids), while triploids represented 75.7%. Both 2:1 *S. cerevisiae:S. paradoxus* (18/33, 54.5%) and 1:2 (7/33, 21.2%) subgenome ratios were observed. However, subgenome balances were not uniform along the genomes of the hybrids, with frequent regions of LOH (**Fig. 1C**). Non-hybrid strains were predominantly diploid, although we identified one tetraploid *S. cerevisiae* and a small fraction of triploid *S. cerevisiae* (10.6%) and *S. paradoxus* (12.5%) isolates.

Variation among the hybrids also extended to their mitochondrial genomes (**Fig. 1D**). The mitochondrial genomes of the hybrids from agave fermentations were frequently recombinant (45.4%), whereas equal proportions were predominantly derived from *S. paradoxus* or *S. cerevisiae* (27.3% each). Thus, the hybrids varied extensively in parental nuclear-genome contribution, ploidy, chromosome-scale allelic composition, and mitochondrial composition (**Fig. 1D**). Together, the geographic breadth and genomic heterogeneity suggest a recurrent *S. cerevisiae* x *S. paradoxus* hybridization system, comprising numerous hybrids and both parental species across multiple agave-spirit producing regions.

### 2. Targeted field sampling reveals a strong association of hybrids with distillery environments

Given that hybridization requires contact between the parental species, we analyzed the evidence of hybrids and parental species co-occurrence within distilleries. We found such co-occurrence in 8 distilleries (12.6%) in eight individual tank samples (6.8%), where either both parental species or one parental species were isolated together with a hybrid (*S. cerevisiae* and *S. paradoxus* n = 2; hybrid and *S. cerevisiae* n = 5; hybrid and *S. paradoxus* n = 1). This local co-occurrence indicated that agave fermentations provide opportunities for interaction between the parental species and for introgression. However, because our initial survey was restricted to distilleries, we could not determine whether hybrids were specifically enriched in fermentation environments or simply reflected a broader regional distribution of their parental species. To distinguish these possibilities, and to better understand the ecological context of the hybrids and their parental populations, we performed three targeted field samplings that included natural areas and agave-spirit distilleries on three regions with distinct hybrid abundance and *S. cerevisiae* diversity (**Fig. 2A**). We selected West I, with the highest proportion of hybrids; South Central, where no hybrids were detected; and Northeast, the only region harboring Mexican Agave 1, the *S. cerevisiae* population with the most introgressions in the Neotropics ^10^. The most distant sampled locations were 1,137 km apart and the elevation range covered from 346 to 2,857 meters above sea level (**Supplementary Data 3**). In these three regions, we visited a total of 15 distilleries where samples were taken from active fermentations, trees and insects, as well as production-associated materials, including tools, surfaces, firewood, and agave residues (all of which we refer to as “distillery materials”). We also sampled 25 sites in natural areas near the distilleries, where bark from trees, insects and fruits were collected (**Fig. 2B, C**; **Methods**).

**Figure 2.**
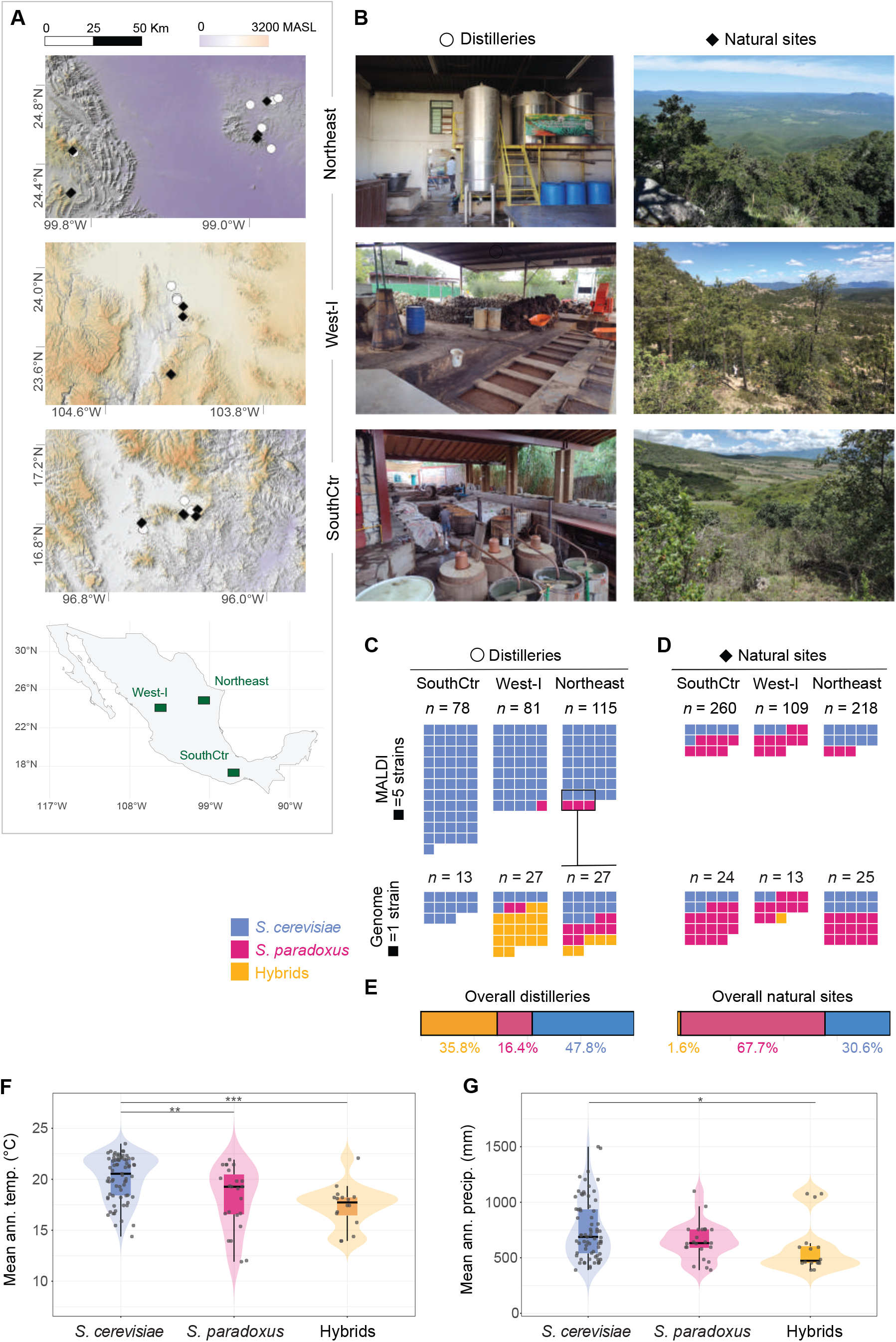
Targeted field sampling reveals a strong association of hybrids with distilleries in contrast to natural environments. A) Geographic location of the three regions where targeted field sampling was performed in distilleries and surrounding natural sites, corresponding to the Northeast, West I and South Central (SouthCtr) agave-spirit producing regions and detailed topographic maps of the sampled sites in the three regions. Sampled distilleries and natural sites are depicted as white circles and black diamonds, respectively. B) Representative images of distilleries and natural sites sampled in each region. C, D) The upper panel shows the isolation rates of *S. cerevisiae* and *S. paradoxus* in distilleries (C) and natural sites (D) across the three sampled regions. The middle panel shows the total numbers of strains assigned to each species by MALDI-TOF biotyping (MALDI) and the bottom panel the numbers of parental and hybrid strains identified by whole-genome sequencing (Genome). In the MALDI panel, each square represents five strains, whereas in the genomic panel, each square represents one strain. E) Aggregated frequency of hybrids, *S. cerevisiae* and *S. paradoxus* identified by whole genome sequencing compared between distilleries and natural sites. F, G) Mean annual temperature (Kruskal−Wallis *p* = 5.0 x 10^−05^, BH−adjusted *p* = 5.0 x 10^−04^) and mean annual precipitation (Kruskal−Wallis *p* = 1.5 x 10^−02^, BH−adjusted *p* = 3.0 x 10^−02^), respectively, of the locations where each strain was isolated. Paired tests BH adjusted are shown as \**p* < 0.05, \*\**p* < 0.01, \*\**p* < 0.001, *S. cerevisiae* n = 71 *S. paradoxus* n = 23, Hybrids n = 16.

We obtained a total of 861 samples from which we generated a collection of 4,006 isolates using enrichment conditions selective for *Saccharomyces*, as we had previously done^34^. Of these, 2,903 isolates (72.5%) could be assigned to a species by MALDI-TOF biotyping, and 2,209 (55.1% of the total collection) met an identification cutoff (score >1.5), including 979 *Saccharomyces* isolates: 843 *S. cerevisiae* and 136 *S. paradoxus*. The overall isolation rate of *Saccharomyces* spp. identified by MALDI-TOF was 40% in samples from distilleries and 5% in samples from natural sites. *S. cerevisiae* was strongly enriched in distilleries relative to natural environments (isolation rates of 39.78% and 3.92%, respectively), whereas assignments to *S. paradoxus* were more evenly distributed between the two environments (5.47% and 3.74%, respectively; **Fig. 2D; Supplementary Fig. 1**). The association between MALDI-assigned species and environment was highly significant (*p* = 3.8 x 10^−7^; Fisher’s exact test), revealing a marked ecological asymmetry between the two parental species.

Given that MALDI-TOF does not identify hybrids, we sequenced the genomes of 129 *Saccharomyces* strains isolated from the targeted sampling in the three regions, 67 from distilleries and 62 from natural sites. Genomic sequencing identified 51 *S. cerevisiae*, 53 *S. paradoxus* and 25 hybrid isolates. Hybrid distribution showed a strong habitat bias. Twenty-four of the 25 genomically identified hybrids were recovered from distilleries, whereas only one from a natural site (24 of 67 versus 1 of 62 sequenced isolates; **Fig. 2C-D and E**; Fisher’s exact test, *p* = 2.2 × 10□). Hybrids were mostly isolated from fermentation tanks, but were also recovered from insects, plants, and production-associated materials within the distilleries (**Supplementary Data 4**). As expected, given the overall distribution of hybrid isolates in the country, most hybrids were recovered in the West I region, including the only one recovered from a natural site, from a beetle. However, targeted sampling also recovered hybrids from distilleries in the Northeast (in the state of Nuevo León), where hybrids had not been detected previously. Thus, despite recovery of both parental species from distilleries and natural environments, hybrids were found almost exclusively in distilleries.

Given the environmental breadth of the targeted sampling, we asked whether hybrid occurrence was associated with climatic and geographic characteristics of the sampling sites (**Fig. 2F,G**). To do so, we considered the newly genomically identified 129 isolates from the targeted sampling together with the 191 from the nationwide agave-fermentation analysis described in the previous section. Hybrids and *S. paradoxus* were associated with localities exhibiting lower mean annual temperatures than those where *S. cerevisiae* was more frequent (**Fig. 2F**). Hybrids were also recovered at higher elevations than *S. cerevisiae*. Similarly, hybrids were found in drier locations with significantly lower mean annual precipitation than *S. cerevisiae* (**Fig. 2G**). Taken together, these observations were reflected in the more frequent occurrence of hybrids in temperate (Cw) and arid to semi-arid (BS) climates. Thus, beyond their strong association with distilleries, hybrid occurrence was geographically structured across the broad range of environmental conditions encompassed by the sampled agave-spirit producing regions.

Together, our targeted sampling revealed a striking ecological asymmetry in which *S. cerevisiae* was strongly enriched in distilleries, *S. paradoxus* was more frequently isolated from natural environments, but also present in anthropogenic environments, and hybrids occurred almost exclusively within distilleries. This pronounced habitat bias extended across environmentally diverse sites, with hybrid occurrence varying along gradients of temperature, elevation, and precipitation.

### 3. A human-associated *S. paradoxus* lineage in agave fermentations

The occurrence of *S. paradoxus* in agave distilleries was striking given the contrasting ecological histories of *S. cerevisiae* and *S. paradoxus*. Whereas *S. cerevisiae* is strongly associated with human fermentations, *S. paradoxus* is predominantly found in natural environments and has only occasionally been reported from anthropogenic fermentation settings^26,27,35–41^. This raised two possibilities, that *S. paradoxus* isolates recovered from distilleries represented local natural populations transiently entering these environments or instead belonged to differentiated human-associated lineages. Distinguishing between these scenarios required solving *S. paradoxus* population structure in Mexico, which remains poorly characterized compared with that of *S. cerevisiae* associated with agave fermentations^10^. We therefore phylogenetically analyzed 63 *S. paradoxus* genomes from our collection, together with 71 strains representing the recognized species clades worldwide^28,42–45^ (**Supplementary Data 2**). In the resulting phylogenies *S. paradoxus* isolates from Mexico clustered within the SpA and SpB clades (**Fig. 3A**). As expected, the majority (52 strains, 82.5%) belonged to SpB, which is considered one of the major North American *S. paradoxus* lineages. Of these, 20 (31.7%) strains were isolated from distilleries (13 from active fermentations and seven from other distillery materials) and 32 (50.8 %) from natural sites.

**Figure 3.**
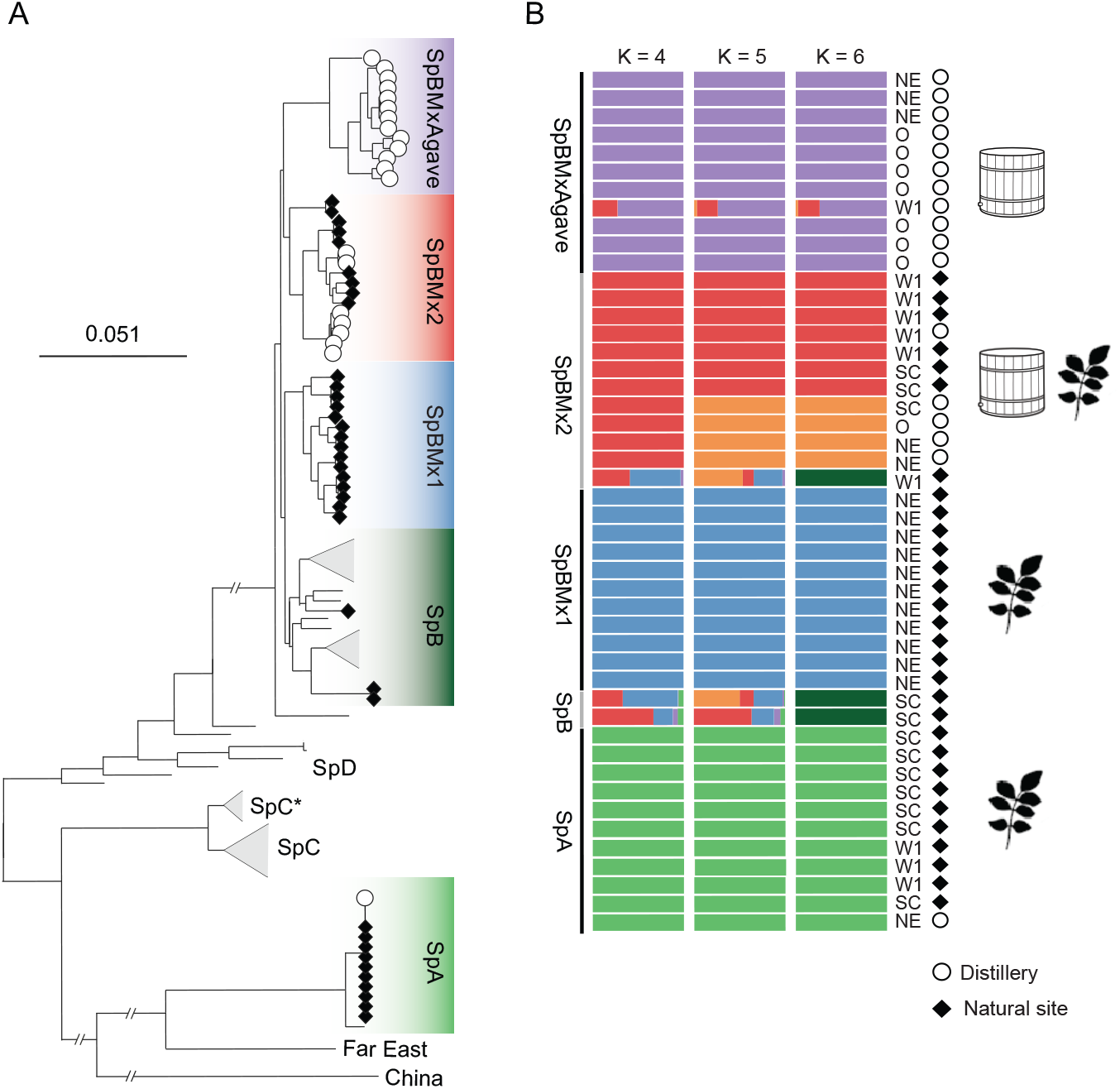
Phylogenetic analysis of *S. paradoxus* strains isolated in Mexico reveal a human-associated lineage. **(A)** Maximum likelihood tree based on 876,358 SNPs from 110 strains. Strains from agave spirit distilleries (21 isolates) and natural sites (37 isolates) are from this study. The known clades of the species are labeled according to previous literature^28,45^, and those containing strains from the current study are colored. At the tip of the branches of the strains identified in the current study the location type is signaled with white circles (○, distilleries) and black diamonds (♦, natural sites). **(B)** Population assignment analysis using ADMIXTURE. At the left of the ADMIXTURE the location type is signaled as in panel A. The agave-spirit producing region is shown at the right side with letters including the three regions of the targeted sampling in this study (Northeast, NE; South Central, SC; and West I, W1) and additional regions from which strains from Gallegos-Casillas *et al.*^34^ were obtained (Others, O). Assignation results for K = 2 to 15 are shown in **Supplementary** Fig. 2.

The phylogeny revealed that the *S. paradoxus* SpB isolates were structured into four subclades (**Fig. 3A**). These subclades are sister to the Brazilian *S. paradoxus* strain formerly known as *S. cariocanus*^46^. One of the subclades from Mexico corresponds to the previously described SpB clade from the U.S.A. and Canada to which natural isolates from the South Central region and West I region were assigned. The remaining three subclades were resolved in this study and are designated here as SpB_Mx lineages. These included (i) SpB_Mx1, a large group of strains from the Northeast region, all isolated from natural sites; (ii) SpB_Mx2, which included strains from both distilleries and natural sites; and (iii) SpB_MxAgave, composed exclusively of strains isolated from distilleries. SpB_Mx1 was closely related to, but clearly distinct from, the North American SpB lineage, forming its sister group (**Fig. 3A**). Subclades SpB_Mx2 and SpB_MxAgave formed a monophyletic group that was more distantly related to the other SpB strains from Mexico, the U.S.A. and Canada.

Population-genomic analyses further supported the differentiation of these Mexican SpB lineages. SpB_MxAgave showed the strongest differentiation from previously characterized North American SpB strains (F_ST_ SpB vs. SpB_MxAgave = 0.49, compared with 0.28 for SpB_Mx1 and 0.29 for SpB_Mx2) and was consistently resolved as a distinct ancestry component in ADMIXTURE analyses across K values (**Fig. 3B; Supplementary Fig. 2**). Evidence of admixture with other populations was detected in only one SpB_MxAgave isolate from West I; the remaining strains showed no detectable admixture. In contrast, SpB_Mx2 strains recovered from distilleries retained ancestry components shared with strains from the surrounding natural environments (**Fig. 3B**).

SpB_MxAgave showed strong ecological association with distilleries: all strains assigned to this lineage were recovered from distillery environments (**Fig. 3A**), and the lineage accounted for 85% (11 of 13) of *S. paradoxus* isolates from active agave fermentations. Yet SpB_MxAgave was geographically widespread, occurring in distilleries across five of the seven agave-spirit-producing regions (Northeast, Northwest, West I, West II, and Balsas Basin), spanning up to 1,440 km. This widespread and repeated occurrence only in human-managed environments across regions was unexpected for a species predominantly associated with wild habitats^26^. Together, these analyses identify SpB_MxAgave as a genetically differentiated and geographically widespread lineage strongly associated with human-managed environments, contrasting with the predominantly wild ecology of *S. paradoxus*.

### 4. Multiple hybrid origins reveal contrasting parental population diversity

To determine whether the agave-associated hybrids originated from a single or multiple ancestral hybridization events, we resolved the parental ancestry of their *S. cerevisiae* and *S. paradoxus* subgenomes. We compared each subgenome with reference panels representing the population diversity of the corresponding parental species (**Methods**). The *S. cerevisiae* reference panel included the diversity previously characterized from agave fermentations in Mexico^10^, natural isolates sequenced here, and representatives of the major global lineages (**Supplementary Fig. 3 and 4**). The *S. paradoxus* panel contained all strains used for the population structure resolved above. For each of the 61 agave-associated hybrids identified through both the initial analysis of previously isolated strains (n = 33), the targeted sampling (n = 25) and other sampling efforts from agave fermentations (n = 3), we independently placed the *S. cerevisiae* and *S. paradoxus* subgenomes within their respective phylogenies and identified their closest non-hybrid relatives (**Fig. 4A,B**). This approach also recapitulated the previously reported parental ancestries of four *S. cerevisiae* x *S. paradoxus* hybrids, isolated from diverse environments, including oak bark in England, a koa tree in Hawaii, a wild habitat in Brazil and an olive-processing environment in Spain^8,9,22,25^.

**Figure 4.**
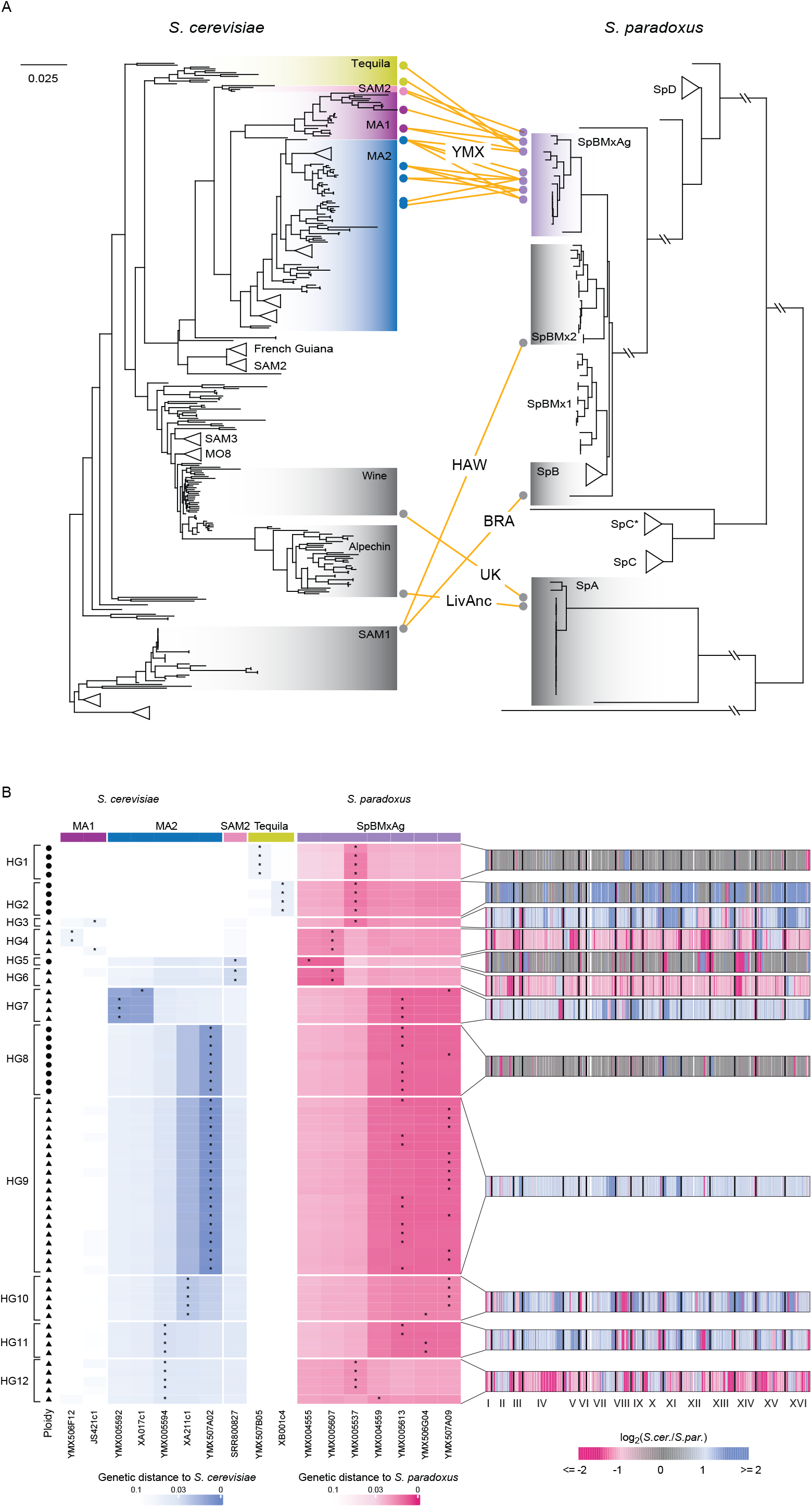
Phylogenetic placement of hybrid subgenomes supports multiple hybridization origins. A) Phylogenetic positions of the *S. cerevisiae* and *S. paradoxus* subgenomes of the hybrids. Orange lines connect the closest *S. cerevisiae* and *S. paradoxus* parental lineages for each unique parental combination identified among the 61 hybrid isolates analyzed in this study and grey lines those of four previously described hybrids. Circles denote strains identified as the closest parental strains of hybrids and are colored according to their clade assignment. Only the parental strains of hybrids identified in this study are shown in color, whereas those of previously identified are shown in grey. B) Phylogenetic-distance profiles of the hybrid subgenomes. Rows represent individual hybrids, and columns represent all non-hybrid *S. cerevisiae* and *S. paradoxus* strains identified as the closest relative of at least one hybrid subgenome. Cell color indicates the genetic distance between each hybrid subgenome and each non-hybrid reference strain, with lower values indicating greater phylogenetic similarity. Asterisks mark the strain or strains with the shortest distance to each subgenome. Symbols on the left indicate ploidy, and brackets indicate the classification into twelve hybrid groups (HG) described in the main text according to their genomic features and parental assignment. Representative chromosome-scale subgenome-composition profiles are shown to the right of the heatmap. Blue indicates genomic regions with a greater contribution from *S. cerevisiae*, magenta indicates homologous regions with a greater contribution from *S. paradoxus*, and grey indicates approximately equal contributions from both subgenomes.

The *S. cerevisiae* subgenomes revealed substantial parental diversity. All 61 agave-associated hybrid subgenomes clustered with Neotropical *S. cerevisiae* strains, with 58 grouping within one of the three main groups identified in Mexico (**Fig. 4A, B**). Most hybrids clustered with the Mexican Agave 2 population (46 hybrids), whereas four grouped with the Mexican Agave 1 and eight with the admixed Tequila Distillery group. The remaining three clustered with the South American Mix 2 (SAM2) clade, which also included an *S. cerevisiae* isolate from a distillery reported here. Across these four populations, ten distinct *S. cerevisiae* strains were identified as the closest sequenced relatives of the hybrid *S. cerevisiae* subgenomes. Thus, the hybrid subgenomes were associated with four genetically differentiated *S. cerevisiae* populations and multiple distinct parental backgrounds, providing strong evidence for recurrent, independent hybridization events.

In contrast, all *S. paradoxus* subgenomes from agave-associated hybrids clustered within a single population, SpB_MxAgave (**Fig. 4A, B)**, despite the presence of at least four other *S. paradoxus* populations in Mexico. The geographic distributions of SpB_MxAgave and hybrids were also concordant. Despite extensive sampling of distilleries and surrounding natural sites in the South Central region, neither was recovered there, although other *S. paradoxus* lineages were present (**Supplementary Fig. 5**). Within SpB_MxAgave, seven distinct strains were identified as the closest sequenced non-hybrid relatives of the 61 *S. paradoxus* hybrid subgenomes. Thus, although the *S. paradoxus* subgenomes all clustered within the same population, their closest relatives included multiple strains within SpB_MxAgave, providing further evidence for independent hybridization events. In contrast, the *S. paradoxus* subgenomes of four previously reported hybrids unrelated to agave fermentations clustered with the SpB and SpB_Mx2 populations, or to the SpA clade. Together, these results revealed a pronounced asymmetry in parental ancestry: the *S. cerevisiae* subgenomes of agave-associated hybrids were associated with several genetically differentiated populations, whereas their *S. paradoxus* subgenomes clustered exclusively with the human-associated SpB_MxAgave population. In fact, nucleotide diversity among the *S. cerevisiae* strains identified as the closest relatives of hybrid subgenomes (π = 0.0036) was higher than that among *S. paradoxus* closest relatives, all belonging to the human-associated SpB_MxAgave clade (π = 0.0012). This asymmetry was also ecological: three of the four contributing *S. cerevisiae* populations occurred in both distillery and natural environments, whereas SpB_MxAgave was restricted to distilleries (**Fig 3A, B; Supplementary Fig. 5**).

We next sought to determine how many independent hybridization events could be distinguished among the agave associated hybrids. Parental subgenome ancestry alone established a minimum of four, corresponding to four distinct combinations of SpB_MxAgave with distinct *S. cerevisiae* populations (**Fig. 4A**). To resolve additional independent events within these combinations, we integrated parental ancestry with subgenome content and ploidy, nuclear heterozygosity patterns, mitochondrial genome type, and, when identifiable, mating-type combination (**Fig. 1**; **Fig. 4B; Supplementary Fig. 6**). Together, these features defined 12 distinct hybrid genomic groups (HG) (**Fig. 4B; Supplementary Fig 7; Table 1**). Some of these groups had similar parental read proportions and ploidy but differed in their closest sequenced relatives and were therefore treated as separate groups. Conversely, groups that differed only in ploidy but shared the same parental population ancestry, similar parental backgrounds, patterns of heterozygosity across the nuclear genome and mitochondrial genome type could represent derivatives of a common ancestral hybridization event. Under this interpretation, HG5 and HG6, and HG8 and HG9, could each derive from a single origin, yielding ten candidate independent events among the 12 HGs. Thus, post-hybridization genome evolution precludes an exact count, but the data supports at least four and up to 12 candidate origins, or ten under the more conservative grouping. Irrespective of the precise number, these analyses reveal recurrent hybridization generating genomically diverse outcomes, while retaining a consistent asymmetry in parental ancestry: multiple *S. cerevisiae* populations contributed to these events, whereas the *S. paradoxus* contribution was restricted to the human-associated SpB_MxAgave population.

**Table 1.** *S. cerevisiae* x *S. paradoxus* hybrid genomic groups identified in agave fermentations.

| **Genomic group<sup>1</sup>** | **No. isolates (61 total)** | **Ploidy** | **Sc lineage<sup>2</sup>** | **Sp lineage<sup>2</sup>** | **Reads mapped to Sc / Sp (% of total)<sup>3</sup>** | **Mt ancestry<sup>4</sup>** | **2µ copy number, range<sup>5</sup>** | **Inferred mating-type combination<sup>6</sup>** | **Geographic distribution<sup>7</sup>** |
|---|---|---|---|---|---|---|---|---|---|
| HG1 | 4 | 2 | TD | SpBMA | 51.8 (0.7) / 45.9 (0.6) | Sc | 40.3–74.9 | Sc-a × Sp-α | CtrHigh |
| HG2 | 4 | 2 | TD | SpBMA | 78.5 (0.6) / 21.0 (0.7) | Sc | 43.7–71.9 | Sc-α × Sp-a | NorthE |
| HG3 | 1 | 3 | MA1 | SpBMA | 68.2 / 31.4 | Sc | 0.0 | N.D. | NorthE |
| HG4 | 3 | 3 | MA1 | SpBMA | 32.7 (0.1) / 66.8 (0.1) | Rec. | 10.6–21.2 | N.D. | CtrHigh (2),<br>Unknown (1) |
| HG5 | 1 | 2 | SAM2 | SpBMA | 47.4 / 52.1 | Rec. | 5.3 | Sc-α × Sp-a | NorthW |
| HG6 | 2 | 3 | SAM2 | SpBMA | 31.9 (2.5) / 67.6 (2.5) | Rec. | 0.0, 10.9 | N.D. | NorthW |
| HG7 | 4 | 3 | MA2 | SpBMA | 70.0 (1.3) / 29.2 (1.3) | Sc | 14.6–25.8 | N.D. | Balsas |
| HG8 | 8 | 2 | MA2 | SpBMA | 48.8 (0.2) / 50.7 (0.3) | Rec. | 0.0–0.0 | Sc-α × Sp-a<sup>(6)</sup> | West1 |
| HG9 | 20 | 3 | MA2 | SpBMA | 64.3 (1.2) / 35.3 (1.0) | Rec. | 3.8–33.0 | N.D. | West1 (19),<br>Balsas (1) |
| HG10 | 5 | 3 | MA2 | SpBMA | 69.1 (0.5) / 30.6 (0.6) | Sc | 0.0–0.0 | N.D. | West1 |
| HG11 | 4 | 3 | MA2 | SpBMA | 60.1 (2.6) / 39.6 (2.5) | Sp | 6.2–20.4 | N.D. | Balsas |
| HG12 | 5 | 3 | MA2 | SpBMA | 34.1 (3.2) / 65.3 (3.1) | Sp | 0.0–0.0 | N.D. | CtrHigh |

<sup>1</sup>Hybrid genomic groups were defined by shared ancestry and genome architecture, as described in Fig. 4B.
<sup>2</sup>Indicates the shared phylogenetic lineage of the closest sampled non-hybrid *S. cerevisiae* relatives within each genomic group. *S. cerevisiae*: TD, Tequila Distillery; MA1, Mexican Agave 1; MA2, Mexican Agave 2; SAM2, South American Mix 2. *S. paradoxus*: SBMA, SpB_MxAgave
<sup>3</sup>Values are mean (SD) percentages of total reads mapped to *S. cerevisiae* / *S. paradoxus*.
<sup>4</sup>Mitochondrial ancestry was assigned from mtDNA read mapping to the parental species; recombinant (Rec.) indicates mitochondrial genomes with mapping patterns inconsistent with a single parental mitotype.
<sup>5</sup>Estimated for each isolate as mean 2µ plasmid coverage divided by median chromosomal coverage; values indicate the range within each genomic group.
<sup>6</sup>Parental mating-type combinations were inferred from species-specific MATa and MATα copy numbers and are reported only for diploid hybrids; triploid isolates were not analyzed (N.D.). For HG8, the inferred combination applies to 7 isolates; one isolate remained unresolved.
<sup>7</sup>Agave-spirit-producing regions in which isolates from each genomic group were recovered.

## Discussion

Across the tree of life, interspecific hybridization is increasingly recognized as an important source of genetic and phenotypic variation^5^, yet the consequences of repeated hybridization remain difficult to resolve in natural populations, particularly when the diversity and ecological distributions of their parental populations are poorly characterized. Here, we uncovered a population-level system of *S. cerevisiae* x *S. paradoxus* hybrids associated with traditional agave fermentations, providing an opportunity to trace recurrent and independent hybridization events in the context of the population diversity of both parental species.

Recurrent interspecific hybridization is not unprecedented in *Saccharomyces*. Multiple independent origins have been inferred for fermentation and natural hybrids^28,47,48^. What distinguishes the agave system is the combination of numerous extant interspecific hybrids, with population-resolved diversity of both parental species, allowing independent hybridization events and their genomic outcomes to be compared within the same system. Our analyses revealed different origins involving distinct parental backgrounds and generating diverse genomic configurations that differed in ploidy, patterns of nuclear heterozygosity, mitochondrial genome type and mating-type combination. This diversity indicates that recurrent hybridization in agave-associated *Saccharomyces* is not restricted to the repeated formation of a single hybrid type but instead produces multiple genomic outcomes. In addition, parental ancestries were consistently asymmetric: several genetically differentiated *S. cerevisiae* populations contributed to the hybrids, while the *S. paradoxus* contribution was restricted to the human-associated SpB_MxAgave population. Together, these findings establish recurrent interspecific hybridization as a population-level feature of agave fermentation-associated *Saccharomyces* and suggest that its occurrence is shaped by both the diversity of available parental backgrounds and lineage-specific factors, rather than simply by the coexistence of the two parental species. The agave fermentation system also challenges the historical ecological contrast between *S. cerevisiae*, whose evolutionary history has been strongly shaped by human-associated environments, and *S. paradoxus*, regarded predominantly as a wild species. Hybrids co-occurred only with the human-associated *S. paradoxus* linage and exclusively in distilleries.

The strong enrichment of hybrids within distilleries raises the question of where hybridization actually occurs and why hybrids accumulate in this environment. Traditional agave fermentations are open systems that continuously exchange microorganisms with insects, plants, and the surrounding landscape, potentially connecting very different stages of the *Saccharomyces* life cycle. Our population-genomic data supports this ecological connectivity: in the South Central and West I regions, *S. cerevisiae* isolates recovered from plants and insects outside distilleries belonged to the same populations as strains from active fermentations and other distillery materials (**Supplementary Fig. 3 and 4**). This pattern was not universal, as the Mexican Agave 1 population was recovered exclusively from distilleries despite extensive sampling of surrounding natural sites, suggesting that the degree of connectivity between natural and human-managed environments differs among *S. cerevisiae* populations. Different stages of the *Saccharomyces* life cycle induced by the open agave fermentation system may be relevant to hybrid formation because active fermentation predominantly favors vegetative growth, whereas nutrient limitation outside tanks could promote sporulation^49^. In the wine system, social-wasp guts can provide successive conditions promoting sporulation, germination, and both intra- and interspecific mating of *Saccharomyces*^50^. Hybrid formation in the agave-fermentation system could similarly occur during environmental phases outside active fermentation, followed by the introduction and proliferation of hybrids within distilleries.

Ecological connectivity alone is unlikely to explain hybrid occurrence. Despite extensive sampling of distilleries and surrounding natural sites, in the South Central region we recovered neither hybrids nor members of SpB_MxAgave although other *S. paradoxus* populations and *S. cerevisiae* were present. Conversely, both hybrids and SpB_MxAgave occurred across multiple other agave-spirit producing regions, especially in the West I, Central Highlands, and Northeast regions, revealing a geographic correspondence between this *S. paradoxus* population and recurrent hybridization. SpB_MxAgave was itself recovered exclusively from distilleries, revealing a human-associated population within a species predominantly characterized from wild habitats. Whether its repeated involvement in hybridization reflects greater opportunities for contact with *S. cerevisiae*, lineage-specific compatibility during hybrid formation, adaptation to distillery conditions, ecological specialization elsewhere in the production landscape, or a combination of these factors remains unsolved.

The genomic diversity observed among agave-associated hybrids arises from two distinct sources: variation already present in the parental strains, which differed among independent hybridization events, and genomic changes accumulating during the subsequent evolution of hybrids. Parental strains can differ in ploidy, heterozygosity if diploid, pre-existing number of introgressions, ancestry composition and mitochrondrial genome type, generating differences among hybrids from the moment they are formed. Hybrid genomes can then diverge further through processes such as LOH, chromosome loss, accumulation of aneuploidies, single nucleotide mutations, and the homogenization of initially biparental mitochondrial genomes. The detailed characterization of the diversity and population structure of both hybridizing species provides the resolution needed to distinguish between these two sources of variation and thereby reconstruct the genomic changes that occurred after independent hybridization events. In this context, the high frequency of triploid hybrids with two copies of one parental subgenome and one copy of the other in agave fermentations is particularly noteworthy. Triploidy could arise through a stepwise route with a cross between diploid and haploid parental cells or saltationally through chromosomal loss following the formation of a tetraploid. Given that the diploids in HG5 and HG8 are very closely related to the triploids in HG6 and HG9, respectively, a stepwise scenario seems more plausible. In these cases, LOH at the mating-type locus could produce a mating competent hybrid that then mated with a haploid from either parental species. This scenario is consistent with recent findings on mechanisms of stepwise triploid formation in *S. cerevisiae*^51^. For the other triploid hybrids lacking a closely related diploid, the same mechanism could have occurred but involving a diploid hybrid that has not been yet identified. Alternatively, a diploid of either parental species could have become mating competent through endoreplication or LOH at the mating-type locus and subsequently mated with a haploid of the other species. Given that 42.6% of the hybrids lack at least one of the four expected regions in the mating-type locus (*Ya* or *Yα* from either *S. cerevisiae* or *S. paradoxus*), the known propensity for recombination and remodeling at the mating-type locus may have contributed to the formation and stabilization of the hybrid lineages.

On the other hand, mitochondrial genome variation of hybrids may provide an example of how post-hybridization processes can reshape initially inherited genomic variation. Newly formed *Saccharomyces* hybrids can initially inherit mtDNA from both parents but rapidly resolve toward one parental mitotype with or without recombination between parental mtDNAs^53–58^. Consistent with this process, agave-associated hybrids carried recombinant mtDNA more frequently than any of the two uniparental forms. Across the 12 hybrid groups, *S. cerevisiae* mitochondrial genomes were the most frequent (five groups), followed by recombinant genomes (five groups), whereas *S. paradoxus* mitochondrial genomes were retained in only two groups. With a single exception, hybrids carrying either *S. cerevisiae* or recombinant mitochondrial genomes also contained a greater proportion of *S. cerevisiae*-derived nuclear material, suggesting some correspondence between mitochondrial genome type and nuclear parental contribution. However, recombinant mitochondrial genomes cannot necessarily be attributed to recombination after hybrid formation, as some of the closest sequenced *S. cerevisiae* relatives of the parental subgenomes already carried recombinant mitochondrial haplotypes. Thus, as with nuclear variation, mitochondrial diversity among the hybrids may reflect both variations inherited from the parental strains already shaped by extensive introgression and genomic changes occurring after hybridization. The functional significance of the observed triploidy and patterns of mitochondrial genome retention remains to be determined.

The long-term evolutionary consequences of hybridization depend on whether initially isolated hybrids can regain fertility and establish gene flow with their parental species. The *S. cerevisiae* Alpechin lineage provides an example of this process: mitotic recombination and progressive LOH can restore regions of sequence homology that facilitate meiotic recombination, providing a route toward fertility, backcrossing and ultimately introgression^9^. Unlike the Alpechin lineage, where the consequences of a single ancestral hybridization were reconstructed through its descendants, the agave fermentation system captures multiple independent hybrid origins at different stages post hybridization. The extensive variation in heterozygosity, ploidy, and parental genome contribution among these hybrids provides a comparative framework for determining whether independent hybrid lineages follow similar trajectories towards fertility and introgression. This question is particularly relevant in the Neotropics, where historical gene flow from *S. paradoxus* has already generated extensive introgression in *S. cerevisiae* populations^8,10,21^. The coexistence of contemporary hybrids with this historical signature of *S. paradoxus* introgression suggests an ongoing cycle of interspecific gene flow in which hybridization is followed by backcrossing and introgression into the parental populations, which may subsequently facilitate further hybridization events.

The combination of multiple independent origins, extant parental populations, and extensive genomic diversity makes *Saccharomyces* from the agave fermentation system particularly powerful for dissecting the processes that shape interspecific hybridization. Independent hybrid lineages provide evolutionary replicates in which the effects of parental background and ecological context can be separated from genomic changes arising after hybridization. Importantly, because *Saccharomyces* is experimentally tractable, these naturally occurring hybrids also provide the opportunity to move directly from population-genomic patterns to controlled tests of the mechanisms that determine genome stabilization, hybrid fitness and fertility. The repeated involvement of the human-associated SpB_MxAgave *S. paradoxus* lineage further provides an opportunity to determine whether ecological context, genetic compatibility, or their interaction contribute to the strongly asymmetric parental contributions observed here. Ultimately, following independent hybrid lineages from their formation through genome stabilization, backcrossing and introgression should reveal why some hybridization events remain transient whereas others establish persistent gene flow and contribute to evolutionary trajectories of their parental species.

## Materials and methods

### Agave fermentation tank sampling

The 191 *Saccharomyces* strains used for the initial identification of hybrids were isolated from fermentation tanks in 63 distilleries (149 strains sequenced in Avelar-Rivas et al. 2026, 43 sequenced in this study), spanning the entire range of regions of Mexico where agave spirits are produced as previously described^34^. Briefly, fermentation samples were collected with sterile serological pipettes from within arm’s reach of the tank edge. Four milliliters were stored in sterile cryovials containing glycerol to a final concentration of 25%. Samples were transported in liquid nitrogen and stored at -70°C until processing.

### Yeast isolation and taxonomic identification

Yeast enrichment was performed using a modified version of the protocol described by Liti *et al.* (2017). Samples were enriched directly in their 50-mL collection tubes by adding 20 mL of *Saccharomyces* sensu stricto enrichment medium and incubating at 25°C for up to three weeks, with periodic inspection for signs of fermentation, including turbidity, sedimentation, and CO_2_ production. Actively fermenting cultures were diluted 10^-3^ – 10^-4^ and plated on WL Nutrient Agar (Sigma and Difco), which facilitates morphological differentiation of yeast colonies. Plates were incubated at 25°C for 2-5 days. For each sample, 12 colonies with *Saccharomyces*-like morphology and 12 colonies representing other morphologies were selected, grown in 150 µL YPD in 96-well plates at 30°C for two days, mixed with glycerol to a final concentration of 20%, and stored at -70°C.

Taxonomic identification was performed by matrix-assisted laser desorption ionization–time of flight mass spectrometry (MALDI-TOF) using a Biotyper® system, following Gallegos-Casillas *et al.*^34^. Isolates were assigned according to similarity scores against the reference database (BDAL v.8 or v.10; scores >1.5). Direct biomass from colonies grown on solid YPD was used for most isolates, with protein extraction performed when necessary to improve spectrum quality.

### DNA extraction and genome sequencing

DNA was extracted using a modified MasterPure^TM^ Yeast DNA Purification Kit protocol. In brief, cells were enzymatically lysed using zymolyase, lyticase, and RNase A, and DNA was precipitated with isopropanol at -20°C. DNA yield was quantified using a Qubit^TM^ fluorometer, and purity was assessed from 260/280 and 260/230 ratios using an ND-1000 NanoDrop^®^. Short-read libraries were prepared and sequenced by BGI on the DNBSeq platform to generate 150-bp paired-end reads.

### Quality control and genome mapping

Raw-read quality was initially assessed by BGI using *SOAPnuke*, and adapters and low-quality reads were removed using *fastp* v0.20.0^59^. Filtered reads were competitively mapped with *bwa* v0.7.4^60^ against a concatenated reference comprising eight *Saccharomyces* species together with *Kluyveromyces marxianus* and *Pichia kudriavzevii*. Reference genomes were *S. cerevisiae* R64-2-1_20150113, *S. paradoxus* YPS138, *S. jurei* GCA_900290405.1, *S. kudriavzevii* GCA_900682695.1, *S. arboricola* GCF_000292725.1, *S. eubayanus* GCA_001298625.1, *S. mikatae* IFO 1815^T^, *S. uvarum* CBS 7001, *K. marxianus* GCF_001417885.1, and *P. kudriavzevii* GCF_003054445.1. The *S. cerevisiae* reference was obtained from SGD, *S. paradoxus* YPS138 from the Yeast Population Reference Panel (YPRP), and the *S. mikatae* and *S. uvarum* assemblies from the Saccharomyces Sensu Stricto resources described by Scannell *et al.*^61^.

### Identification of hybrid isolates

Following competitive mapping to the multispecies concatenated reference, reads were assigned according to the species reference to which they mapped. Isolates with >87.5% of mapped reads assigned to a single species nuclear genome were classified as non-hybrid representatives of that species, whereas isolates with ≥12.5% of mapped reads assigned to two nuclear *Saccharomyces* genomes were classified as hybrids (the expected percentage following two backcrosses of an F1 hybrid to one of the parental lineages).

### Determination of ploidy

Ploidy was determined by flow cytometry analysis of the isolates stained with propidium iodide using lab strains BY4741 and BY4743 as haploid and diploid controls respectively to at least one representative strain of each Hybrid Group. For strains without a cytometry measurement, ploidy was estimated from sequence coverage supporting each different allele at a given position. For hybrid ploidy estimation the parental subgenome with the lowest median coverage was assumed to represent a single copy, and the ploidy of the other parental subgenome was assigned as the ratio of its median coverage to that of the lowest coverage subgenome. The approach was validated with the isolates in which both the cytometry and the coverage-based inference showed full correspondence.

### Determination of nuclear subgenome composition in hybrids

To assess the genomic contribution of each parental species, a chain alignment between the *S. cerevisiae* S288C and *S. paradoxus* YPS138 reference genomes was generated using the doblastzchainnet.pl from the Kent utilities^62,63^, with *S. cerevisiae* as the query and *S. paradoxus* as the target. Positions with multiple alignments, mitochondrial and 2μ plasmid were removed. The filtered chain alignment was used to establish one-to-one correspondence between homologous positions of the two parental reference genomes. For each parental subgenome, sequencing depth was calculated in non-overlapping 1-kb windows defined by *S. cerevisiae* coordinates. Only windows with at least 80% shared sequence homology were considered. Parental genome dosage was then estimated for each window as the log_2_ ratio of *S. cerevisiae* to *S. paradoxus* sequencing depth.

### Determination of mitochondrial ancestry and 2_μ_m plasmid copy number

The concatenated reference included the mitochondrial genomes of all *Saccharomyces* species, allowing mitochondrial ancestry to be inferred directly from competitive mapping. Mitochondrial composition was estimated from the proportion of reads mapping to the *S. cerevisiae* and *S. paradoxus* mitochondrial genomes. Mitotype was considered recombinant when the competitive mapping showed at least 12.5% of mitochondrial reads assigned to the two species. The relative copy number of the *S. cerevisiae* 2μ plasmid was estimated by dividing its median coverage by the lower of the median nuclear coverages of the two parental subgenomes (*S. cerevisiae* or *S. paradoxus*), providing a single-copy nuclear reference regardless of hybrid ploidy.

### Targeted field sampling

Sampling was conducted in three regions: South Central (Oaxaca), West I (Durango), and Northeast (Nuevo León and Tamaulipas). These regions encompass distinct biogeographic provinces, including the *Sierra Madre del Sur*, *Chihuahua Desert*, *Sierra Madre Oriental*, and *Tamaulipas* provinces^64^. In total, 15 distilleries and 25 natural sites were sampled. Natural sites were selected near sampled distilleries but in areas with minimal human disturbance and with *Quercus* trees present. Sites were located 1–42 km from the nearest distillery (mean, 9.2 km), with most at least ∼3 km and up to 42 km away. Samples from fermentation tanks in distilleries were obtained as formerly described^34^.

#### Distillery materials

Within distilleries, parts of agave plants, tools, and production-associated surfaces were sampled, including cooked agave stems, grinding residues, remnants of previous fermentations, firewood, pounders, and wooden or metal tools. Such sampling was also performed in distilleries that were temporarily inactive. Collection instruments were disinfected between samples with 10% chlorine followed by 70% ethanol.

#### Plants

In distilleries, bark was collected from trees growing within or along the boundaries of the distillery premises and having a diameter at breast height (DBH) >10 cm. At natural sites, 50 x 10 m transects were established and bark was sampled from oaks and other plant families occurring within the same transects with DBH >10 cm. Approximately 8–20 g of bark was collected into sterile 50-mL conical tubes using autoclaved cork borers, knives, and tweezers, which were disinfected between samples with 10% chlorine followed by 70% ethanol.

#### Insects

Insects were actively collected using entomological nets UV-sterilized before sampling. Nets were swept approximately 1 m above ground until 10–20 insects were captured and transferred to sterile 50-mL conical tubes using an autoclaved aspirator. Nets were disinfected with 70% ethanol between collections within a site, and a newly sterilized net was used at each new location. All samples from distillery materials, plants and insects were kept on ice for 3–5 days until laboratory processing (enrichment and incubation).

#### Field metadata

Sample metadata including date, location type, location name, geographical coordinates, sample type, photograph, and collector, were recorded using KoboCollect and stored on the KoboToolbox server. Data were cleaned using *Python* v3.8.8 and *OpenRefine* v3.4.1. Overall, 861 samples collected in the field yielded 4,006 yeast isolates, of which 1,094 isolates from 139 samples were identified as *Saccharomyces* by MALDI-TOF. Strains were selected for genome sequencing based on MALDI-TOF identity scores, geographical origin, and isolation substrate. Yeast isolation, DNA extraction, sequencing, quality control and competitive read mapping were performed with collected samples as described above.

### Environmental data

Environmental characteristics of the collection sites were obtained by mapping the location points (as from their geographical coordinates recorded in the field) over the corresponding cartography. Elevation data was obtained at 1 km scale resolution through the US Geological Survey GTOPO30 dataset^65^. WorldClim climatic layers were used to obtain Mean Annual Precipitation (wcbio_12) and Mean Annual Temperature (wcbio_01) at 1 km (30 arc-seconds) resolution^66^. Climate types were determined using the Köppen classification system modified by García^67^.

### Variant calling, genotyping, and phylogenetic analysis

All isolates classified as *S. paradoxus* by competitive genome mapping were remapped to the YPS138 reference genome. Variant calling was performed with GATK v4.1.1.0^68^, allowing up to 10% of missing SNPs per sample. The final *S. paradoxus* dataset comprised 110 genomes and 876,358 SNPs. The dataset included 21 strains from distilleries and 37 from natural sites (all from this study), as well as 52 strains representative of the recognized clades of the species SpA, SpB, SpC, SpC*, SpD, Far East, and China (references in **Supplementary Data 2**).

An equivalent pipeline was applied to isolates identified as *S. cerevisiae*, which were mapped to the S288C reference genome. The dataset additionally included 47 strains from agave fermentations from other regions in Mexico, including seven strains originally isolated by Lachance^69^, and 105 strains representing major worldwide lineages (references in **Supplementary Data 2**). The final *S. cerevisiae* dataset comprised 203 genomes and 870,190 biallelic SNPs.

Each species’ matrix with the genotyped SNPs was aligned and converted from the default GATK vcf format to PHYLIP format with a python script^70^. Afterwards, the phylogenetic trees were built using the maximum likelihood algorithm of *RAxML* v8.2.12 and a GTR substitution model^71^. We used 100 bootstraps to assess branch support.

### Population structure analysis

Population structure was assessed using *ADMIXTURE* v1.3.0^72^ separately for strains of each species isolated from the three regions of the targeted sampling. To avoid any bias in the analysis related to possible clonal replicates, we removed strains that were isolated from the same sample and that displayed a phylogenetic distance of < 0.005 in the ML phylogenetic tree. In those cases, we chose at random one strain from the group of isolates suspected to be clones. The *S. paradoxus* analysis included 47 strains, all from this study. The *S. cerevisiae* analysis included 88 strains, comprising 36 from this study, 46 from Avelar-Rivas *et al.*^10^, and 6 from the Mexican Agave 1 lineage reported by Peter *et al.*^21^. For each species, the corresponding SNP matrix was restricted to these samples and pruned for linkage disequilibrium using *plink* v1.9 (--indep-pairwise 50 5 0.5). The resulting datasets contained 108,636 SNPs for *S. cerevisiae* and 86,448 for *S. paradoxus*. *ADMIXTURE* was run for K = 1-15 using 10 independent seeds per each K, and the optimal K was selected based on the lowest cross-validation error (**Supplementary Figs. 2 and 4**).

### Phylogenetic placement of hybrid subgenomes

A set of 65 *S. cerevisiae* x *S. paradoxus* hybrids was analyzed, comprising 61 hybrid strains isolated from Mexico, the Living Ancestor from the Alpechin clade^9^ and one isolate each from the United Kingdom, Hawaii, and Brazil^8,25^. Raw reads were processed and mapped to the multispecies concatenated reference as described above, and variants were called using GATK v4.1.1^68,73^, allowing 10% of missing SNPs per sample. To prevent competitive mapping bias in the hybrid and *S. paradoxus* samples due to the lack of ribosomal DNA (rDNA) sequences in the *S. paradoxus* YPS138 reference genome, reads mapping to the rDNA sequence coordinates of the *S. cerevisiae* S288C reference were masked in all samples. Hybrid variants were then separated into *S. cerevisiae* and *S. paradoxus* nuclear subgenome-specific files.

For phylogenetic placement, each hybrid subgenome was analyzed independently together with a population-resolved reference panel of the corresponding parental species. The *S. cerevisiae* reference panel comprised 512 strains, including 20 strains from the Mexican Agave 1 lineage, 176 strains from the Mexican Agave 2 lineage, and 16 strains from the Tequila Distillery lineage, along with representatives of major worldwide lineages (**Supplementary Data 2**). The *S. paradoxus* reference panel comprised 134 strains, including 11 strains from the SpB_Mx_Agave lineage, 12 from the SpB_Mx1 lineage, and 12 from the SpB_Mx2 lineage, as well as the Mexican representatives of SpB and SpA, and strains encompassing the diversity of the major global clades (**Supplementary Data 2**).

For each hybrid independently, its subgenome GVCF was jointly genotyped with the corresponding parental reference panel using GenotypeGVCFs. Variants were restricted to SNPs and quality-filtered following GATK best practices, removing sites with >10% missing data with *vcftools*^74^. Alignments were converted to PHYLIP format^70^, and sites lacking information in the hybrid itself were removed so that all retained positions contributed to its phylogenetic placement. Maximum-likelihood phylogenies were reconstructed with RAxML v8.2.12 under the GTRGAMMA model^71^, with 100 rapid-bootstrap replicates for *S. paradoxus* and 10 for *S. cerevisiae*. Each *S. cerevisiae* phylogenetic placement analysis contained 513 strains (512 of the reference panel plus one hybrid subgenome) and an average of 1,039,415 SNPs (SD = 138,724; *n*= 65 phylogenetic trees), whereas each *S. paradoxus* analysis contained 135 strains (134 from the reference panel plus one hybrid subgenome) and an average of 874,039 SNPs (SD = 172,263; *n* = 65 phylogenetic trees).

### Assignment of closest strain to hybrid subgenomes

For each parental species (*S. cerevisiae* and *S. paradoxus*), the closest reference strain to each hybrid subgenome was identified from the corresponding hybrid-specific maximum-likelihood phylogeny. Pairwise phylogenetic distances, calculated as the sum of branch lengths separating the hybrid from each non-hybrid reference strain, were obtained using the Biopython Phylo module^75^. The non-hybrid strain with the minimum distance to each hybrid subgenome was designated as the closest reference strain. The closest reference strain represents the most similar available sequence for each parental species, related to but not necessarily the progenitor of the hybrid.

### Copy number of the mating-loci Y-regions in both species

To estimate the copy number of the Ya and Yα regions in the diploid hybrid genomes, coverage depth was extracted from alignments without mapping-quality filtering, at the corresponding Y region of the three *HML*, *MAT* and *HMR* loci in each parental subgenome. For Yα, present in both *HML* and *MAT* loci of the two reference genomes, coverage depth was summed across these two loci for each subgenome. For Ya that is only present at the *HMR* locus, coverage depth was taken only from this locus. The resulting values for Ya-*S. cerevisiae*, Ya-*S. paradoxus*, Yα-*S. cerevisiae* and Yα-*S. paradoxus* were each divided by the median nuclear-genome-wide coverage of the parental subgenome with the lower median coverage and rounded to the nearest integer to estimate the copy number of each region.

## Data accessibility

Genome sequencing data generated in this study have been deposited in the NCBI SRA under the BioProject accession <u>PRJNA1232640</u>. The accession numbers of each genome employed in this study, including those previously sequenced, are provided in **Supplementary Data 2**. A Spanish translation of the article is provided in **Supplementary Data 5**.

## Authorship contributions

Conceptualization: J.A.A-R., X.A-D., A.D.L., L.M. and E.M.; methodology: J.A.A-R., C.L-G., I.S., and L.M.; software: J.A.A-R., I.S., R.E-S. and E.S-G; validation: J.A.A-R. and I.S.; formal analysis: J.A.A-R., X.A-D., C.L-G., I.S., R.E-S., E.S-G. and M.A.; investigation: J.A.A-R., X.A-D., C.L-G., I.S., R.R-S., E.S-G. and M.A.; resources: M.R.K., G.L., A.D.L., L.M. and E.M.; data curation: J.A.A-R., X.A-D., C.L-G., I.S., R.E-S., E.S-G and L.M.; writing-original draft: J.A.A-R., X.A-D., C.L-G., A.D.L., L.M. and E.M.; writing-review and editing: I.S. and G.L.; visualization: J.A.A-R., X.A-D., C.L-G., I.S., R.E-S., A.D.L. and E.M.; supervision: M.R.K., G.L., A.D.L., L.M. and E.M.; project administration: A.D.L., L.M. and E.M.; funding acquisition: G.L., A.D.L., L.M. and E.M. All authors read and approved the final version of the manuscript.

## Declaration of competing interest

The authors declare no competing interests. Two preprints are associated to this work: https://doi.org/10.1101/2025.05.31.656962 and https://doi.org/10.64898/2026.11.04.764348.

## Acknowledgements

We are grateful to the producers and everyone involved in agave spirit production who granted us access to their distilleries and supported us during our fieldwork, particularly Oscar Estrada, Gerardo Hernández, José Juan Simental, Ramiro Ávila, Gilberto Roldán, Cuauhtémoc Jacques, Alejandro González, Karina Abad, Graciela Ángeles and Edgar Ángeles. We thank Luis Aguilar, Vanessa Arellano, Antonio Basurto, Marco Chavez, Mauricio Campa, Alejandra Castillo, Maritrini Colón, Aaron de Luna, Porfirio Gallegos, Jair García, Luis F. García, Antonio Hernández, Christian Molina, Martín Moreno, Susana Ruiz, José Luis Téllez, Carina Uribe, and Karen Vaca for technical assistance, and Anne Gschaedler, Marc Andre Lachance and Luis Delaye for sharing strains and resources.

## Funding

This work was funded by Secretaría de Ciencia, Humanidades, Tecnología e Innovación (Secihti) grants CF-2023-G-695, CBF-2025-G-838, and CBF-2026-3450, and by UNAM-PAPIIT grant IN212524. G.L. and J.A.A-R. were supported by the Impulscience - Fondation Bettencourt Schueller

## Declaration of generative AI and AI-assisted technologies in the manuscript preparation process

During the preparation of this work, the authors used ChatGPT, Claude and DeepSeek to assist with the development, modification, adaptation, and debugging of existing scripts used for data processing and visualization, as well as with English-language editing to improve clarity and readability. The authors reviewed, tested, and validated the generated code and its outputs, and take full responsibility for the analyses, interpretation of the results, and content of the published article.

## Supplementary material

**Supplementary Data 1**. List of the strains sequenced in this study, with associated sampling and genomic metadata.

**Supplementary Data 2**. List of strains and associated metadata used in this study.

**Supplementary Data 3**. List of sampled natural sites and distilleries, and their associated metadata.

**Supplementary Data 4**. List of hybrids and associated metadata used in this study.

**Supplementary Data 5**. Spanish translation of the article.

**Supplementary Figure 1.** Isolation of *Saccharomyces* spp. from the three geographical areas of Mexico where targeted sampling was performed. The bar plot shows the isolation rates as the percentage of samples that yielded *S. cerevisiae* (grey bars) or *S. paradoxus* (black bars) confidently identified with MALDI-TOF (score > 1.5) from each substrate type in distilleries and natural sites. Sample size of each category is shown on top.

**Supplementary Figure 2.** Extended ADMIXTURE analysis of the *S. paradoxus* strains identified in the targeted sampling. A) ADMIXTURE plots for 47 *S. paradoxus* strains from *K* = 2 to *K* = 15, with 86,448 SNPs (previously LD pruned). B) ADMIXTURE cross-validation errors for *K* = 2 to *K* = 15, in a matrix of 86,448 SNPs (previously LD pruned) and 47 *S. paradoxus* strains. The red line represents the mean, whereas the green line represents the median.

**Supplementary Figure 3.** Relationships of *S. cerevisiae* strains from agave spirit distilleries and natural sites in Mexico. **(A)** Maximum likelihood tree using 870,190 biallelic SNPs of 203 strains. Strains from agave spirit distilleries include 42 isolates from this study, 56 from ^10^, and 8 from ^21^. All 19 *S. cerevisiae* isolates from natural sites in Mexico are from this study. The main clades of the world are indicated following the nomenclature of Peter and DeChiara, *et al.*^21^ (with the exception of Wild Brazil 1 and 3 from^8^ and Pacific West Coast from^76^); the Neotropical cluster was described by Avelar-Rivas *et al.*^10^. Only clades that include strains from this study or Avelar-Rivas *et al.*, 2026 are colored. Agave spirit producing regions only include the three regions visited during the targeted sampling in this study (Northeast, South Central and West I). Location type is indicated with white circles (○, distilleries) or black diamonds (♦, natural sites) at the tip of each branch**. (B)** Population assignment analysis using ADMIXTURE. Region is shown at the right; location type is indicated with white circles for distilleries and black diamonds for natural sites at the left. Assignation results for K = 2 to 15 are shown in **Supplementary Figure 4**.

**Supplementary Figure 4.** ADMIXTURE analyses of *S. cerevisiae* strains from agave-spirits distilleries and natural sites of Mexico collected during the targeted sampling effort. **(A)** ADMIXTURE plots of 88 *S. cerevisiae* strains from K = 2 to K = 15. **(B)** ADMIXTURE cross-validation errors for K=2 to K=15, in a matrix of 108,636 SNPs (previously LD pruned) and 88 *S. cerevisiae* strains. The red line represents the mean, whereas the green line represents the median.

**Supplementary Figure 5.** Geographic distribution of *S. cerevisiae* and *S. paradoxus* populations associated with agave fermentations and natural sites in Mexico. Circular insets summarize the occurrence of *S. cerevisiae* (circles) and *S. paradoxus* (squares) subpopulations in each region sampled (Central Highlands [CtrHigh]; South Central [SouthCtr]; Balsas Basin [BalsasB]). Colors indicate the subpopulation assignments shown in the schematic trees at left. Symbols located inside the white circles represent strains isolated from distilleries, whereas symbols outside the circles correspond to strains recovered from natural sites that were sampled only in the West I, Northeast, and South Central regions. Symbol size is scaled according to the number of isolates belonging to each population within a region, with the visual scale capped at *n*=30; groups with more than 30 isolates are therefore shown at the maximum symbol size. Actual isolate counts range from 1 to 76. Red connecting solid lines indicate the occurrence of interspecific hybrids between *S. cerevisiae* and *S. paradoxus* subpopulations. Line thickness is scaled according to the number of independent hybrid strains inferred for each parental combination, with the visual scale capped at *n*=30; combinations with more than 30 hybrids are therefore shown at the maximum line width. Actual hybrid counts range from 1 to 32. Empty symbols connected by yellow lines indicate inferred parental populations for which no non-hybrid parental isolate was recovered from the corresponding region.

**Supplementary Figure 6.** Mating-type loci configurations in diploid Hybrid Groups support independent hybridization events. The first column shows the per-base coverage across the mating-type-specific *Ya* and *Yα* regions of both parental species. No mapping-quality filter was applied for this analysis allowing multimapping reads with equally good alignments to be randomly assigned to one of the equally-scoring positions. The coverage of the two identical *Yα* regions was added. Boxplots show the median and interquartile range, with whiskers extending 1.5x the ICR. In column two, copy-number estimates show relative copy numbers of *Ya* and *Yα* regions in each species. This was calculated by dividing each median by the smallest non-zero median coverage within each isolate and rounding to the nearest integer. The third column shows the mean 1kb windowed coverage after filtering as described in Methods across Chromosome III. The *HML*, *MAT* and *HMR* loci are shown, highlighting the location of the *Ya* or *Yα* in both subgenomes in the concatenated reference. Coverage at the mating-type cassettes themselves is depleted at Q30 filtering because reads from these homologous regions frequently map ambiguously; therefore, local ancestry was inferred from the pattern of allelic balance at the flanking regions. The last column shows the inferred mating-type locus configuration, representing the most parsimonious arrangement consistent with the estimated *Y*-region copy numbers and the chromosome III ancestry together with LOH patterns shown in column three.

**Supplementary Figure 7.** Relative subgenome coverage of the 61 hybrids sequenced in this study. The log_2_ ratio of *S. cerevisiae* to *S. paradoxus* sequencing depth, calculated in 1-kb windows with at least 80% of bases aligned between the parental reference genomes is used as a proxy for subgenome balance. The resulting profiles reveal distinct patterns among hybrid groups (HGs), as well as variation within individual groups. Hybrids are ordered as in Fig. 4B, with members of each HG displayed consecutively.

## Notes

### Competing Interest Statement

The authors have declared no competing interest.

### Summary of Updates

The author order was corrected since it did not match the order in the uploaded manuscript. In addition, we made corrections to the main text.

## References

1. Stelkens, R. B., Brockhurst, M. A., Hurst, G. D. D., Miller, E. L. & Greig, D. The effect of hybrid transgression on environmental tolerance in experimental yeast crosses. j. evol. Biol. 27, 2507–2519 (2014).

2. Samarasinghe, H., You, M., Jenkinson, T. S., Xu, J. & James, T. Y. Hybridization Facilitates Adaptive Evolution in Two Major Fungal Pathogens. Genes 11, 101 (2020).

3. Steensels, J., Gallone, B. & Verstrepen, K. J. Interspecific hybridization as a driver of fungal evolution and adaptation. Nat Rev Microbiol 19, 485–500 (2021).

4. Thompson, K. A. et al. The Ecology of Hybrid Incompatibilities. Cold Spring Harb Perspect Biol 16, a041440 (2024).

5. Schumer, M. & Rieseberg, L. H. Genetics and genomics of hybridization. Nat Rev Genet 1– 15 (2026) doi:10.1038/s41576-026-00995-7.

6. Arnold, M. L. & Martin, N. H. Adaptation by introgression. J Biol 8, 82 (2009).

7. Morales, L. & Dujon, B. Evolutionary Role of Interspecies Hybridization and Genetic Exchanges in Yeasts. Microbiology and Molecular Biology Reviews 76, 721–739 (2012).

8. Barbosa, R. et al. Evidence of Natural Hybridization in Brazilian Wild Lineages of Saccharomyces cerevisiae. Genome Biol Evol 8, 317–329 (2016).

9. D’Angiolo, M. et al. A yeast living ancestor reveals the origin of genomic introgressions. Nature 587, 420–425 (2020).

10. Avelar-Rivas, J. A. et al. Recurrent introgression and geographical stratification shape Saccharomyces cerevisiae in the Neotropics. Nat Commun 17, 3024 (2026).

11. Denef, V. J. & Banfield, J. F. In Situ Evolutionary Rate Measurements Show Ecological Success of Recently Emerged Bacterial Hybrids. Science 336, 462–466 (2012).

12. Edelman, N. B. & Mallet, J. Prevalence and Adaptive Impact of Introgression. Annual Review of Genetics 55, 265–283 (2021).

13. Rosser, N. et al. Hybrid speciation driven by multilocus introgression of ecological traits. Nature 628, 811–817 (2024).

14. Frayer, M. E., Robles, N. V., Rodríguez-Barrera, M. J., Coughlan, J. M. & Schumer, M. The molecular evolutionary basis of species formation revisited. Trends in Genetics 41, 1068– 1095 (2025).

15. Delneri, D. et al. Engineering evolution to study speciation in yeasts. Nature 422, 68–72 (2003).

16. Huerta-Sánchez, E. et al. Altitude adaptation in Tibetans caused by introgression of Denisovan-like DNA. Nature 512, 194–197 (2014).

17. Sankararaman, S., Mallick, S., Patterson, N. & Reich, D. The Combined Landscape of Denisovan and Neanderthal Ancestry in Present-Day Humans. Current Biology 26, 1241– 1247 (2016).

18. Harris, D. N. et al. Diverse African genomes reveal selection on ancient modern human introgressions in Neanderthals. Current Biology 33, 4905–4916.e5 (2023).

19. Arnold, M. L. & Hodges, S. A. Are natural hybrids fit or unfit relative to their parents? Trends in Ecology & Evolution 10, 67–71 (1995).

20. Mallet, J. Hybridization as an invasion of the genome. Trends in Ecology & Evolution 20, 229–237 (2005).

21. Peter, J. et al. Genome evolution across 1,011 Saccharomyces cerevisiae isolates. Nature 556, 339–344 (2018).

22. Pontes, A., Čadež, N., Gonçalves, P. & Sampaio, J. P. A Quasi-Domesticate Relic Hybrid Population of Saccharomyces cerevisiae × S. paradoxus Adapted to Olive Brine. Front. Genet. 10, (2019).

23. Peris, D. et al. Macroevolutionary diversity of traits and genomes in the model yeast genus Saccharomyces. Nat Commun 14, 690 (2023).

24. Loegler, V., Friedrich, A. & Schacherer, J. Overview of the Saccharomyces cerevisiae population structure through the lens of 3,034 genomes. 2024.09.16.613241 Preprint at 10.1101/2024.09.16.613241 (2024).

25. Tellini, N. et al. Ancient and recent origins of shared polymorphisms in yeast. Nat Ecol Evol 8, 761–776 (2024).

26. Boynton, P. J. & Greig, D. The ecology and evolution of non-domesticated Saccharomyces species. Yeast 31, 449–462 (2014).

27. Nikulin, J. et al. Brewing potential of the wild yeast species Saccharomyces paradoxus. Eur Food Res Technol 246, 2283–2297 (2020).

28. Eberlein, C. et al. Hybridization is a recurrent evolutionary stimulus in wild yeast speciation. Nat Commun 10, 923 (2019).

29. Marcet-Houben, M. & Gabaldón, T. Beyond the Whole-Genome Duplication: Phylogenetic Evidence for an Ancient Interspecies Hybridization in the Baker’s Yeast Lineage. PLOS Biology 13, e1002220 (2015).

30. Bendixsen, D. P., Frazão, J. G. & Stelkens, R. Saccharomyces yeast hybrids on the rise. Yeast 39, 40–54 (2022).

31. Gallone, B. et al. Interspecific hybridization facilitates niche adaptation in beer yeast. Nat Ecol Evol 3, 1562–1575 (2019).

32. Dunn, B. & Sherlock, G. Reconstruction of the genome origins and evolution of the hybrid lager yeast Saccharomyces pastorianus. Genome Research 18, 1610–1623 (2008).

33. Libkind, D. et al. Microbe domestication and the identification of the wild genetic stock of lager-brewing yeast. Proceedings of the National Academy of Sciences 108, 14539–14544 (2011).

34. Gallegos-Casillas, P. et al. Yeast diversity in open agave fermentations across Mexico. Yeast 41, 35–51 (2024).

35. Naumov, G. I., Naumova, E. S. & Sniegowski, P. D. Saccharomyces paradoxus and Saccharomyces cerevisiae are associated with exudates of North American oaks. Can. J. Microbiol. 44, 1045–1050 (1998).

36. Sniegowski, P. D., Dombrowski, P. G. & Fingerman, E. Saccharomyces cerevisiae and Saccharomyces paradoxus coexist in a natural woodland site in North America and display different levels of reproductive isolation from European conspecifics. FEMS Yeast Res 1, 299–306 (2002).

37. Sweeney, J., Kuehne, H. & Sniegowski, P. Sympatric natural and populations have different thermal growth profiles. FEMS Yeast Research 4, 521–525 (2004).

38. Sampaio, J. P. & Gonçalves, P. Natural Populations of Saccharomyces kudriavzevii in Portugal Are Associated with Oak Bark and Are Sympatric with S. cerevisiae and S. paradoxus. Applied and Environmental Microbiology 74, 2144–2152 (2008).

39. Zhang, H., Skelton, A., Gardner, R. C. & Goddard, M. R. Saccharomyces paradoxus and Saccharomyces cerevisiae reside on oak trees in New Zealand: evidence for migration from Europe and interspecies hybrids. FEMS Yeast Res 10, 941–947 (2010).

40. Dashko, S. et al. Changes in the Relative Abundance of Two Saccharomyces Species from Oak Forests to Wine Fermentations. Front. Microbiol. 7, (2016).

41. Boynton, P. J., Kowallik, V., Landermann, D. & Stukenbrock, E. H. Quantifying the efficiency and biases of forest Saccharomyces sampling strategies. Yeast 36, 657–668 (2019).

42. Leducq, J.-B. et al. Local climatic adaptation in a widespread microorganism. Proc. R. Soc. B 281, 20132472 (2014).

43. Xia, W. et al. Population genomics reveals structure at the individual, host-tree scale and persistence of genotypic variants of the undomesticated yeast Saccharomyces paradoxus in a natural woodland. Molecular Ecology 26, 995–1007 (2017).

44. Yue, J.-X. et al. Contrasting evolutionary genome dynamics between domesticated and wild yeasts. Nat Genet 49, 913–924 (2017).

45. He, P.-Y. et al. Highly diverged lineages of Saccharomyces paradoxus in temperate to subtropical climate zones in China. Yeast 39, 69–82 (2022).

46. Liti, G., Barton, D. B. H. & Louis, E. J. Sequence Diversity, Reproductive Isolation and Species Concepts in Saccharomyces. Genetics 174, 839–850 (2006).

47. Peris, D., Lopes, C. A., Arias, A. & Barrio, E. Reconstruction of the Evolutionary History of Saccharomyces cerevisiae x S. kudriavzevii Hybrids Based on Multilocus Sequence Analysis. PLOS ONE 7, e45527 (2012).

48. Baker, E. et al. The Genome Sequence of Saccharomyces eubayanus and the Domestication of Lager-Brewing Yeasts. Mol Biol Evol 32, 2818–2831 (2015).

49. Freese, E. B., Chu, M. & Freese, E. Initiation of Yeast Sporulation by Partial Carbon, Nitrogen, or Phosphate Deprivation. JOURNAL OF BACTERIOLOGY https://doi.org/10.1128/jb.149.3.840-851.1982 (1982) doi:10.1128/jb.149.3.840-851.1982.

50. Stefanini, I. et al. Social wasps are a Saccharomyces mating nest. Proceedings of the National Academy of Sciences 113, 2247–2251 (2016).

51. Gómez-Muñoz, C. et al. A postmeiotic route to stepwise polyploidy. 2026.02.20.706944 Preprint at 10.64898/2026.02.20.706944 (2026).

52. Mozzachiodi, S. et al. Aborting meiosis allows recombination in sterile diploid yeast hybrids. Nat Commun 12, 6564 (2021).

53. Antunovics, Z., Nguyen, H.-V., Gaillardin, C. & Sipiczki, M. Gradual genome stabilisation by progressive reduction of the Saccharomyces uvarum genome in an interspecific hybrid with Saccharomyces cerevisiae. FEMS Yeast Res 5, 1141–1150 (2005).

54. Leducq, J.-B. et al. Mitochondrial Recombination and Introgression during Speciation by Hybridization. Mol Biol Evol 34, 1947–1959 (2017).

55. Verspohl, A., Pignedoli, S. & Giudici, P. The inheritance of mitochondrial DNA in interspecific Saccharomyces hybrids and their properties in winemaking. Yeast 35, 173–187 (2018).

56. Hewitt, S. K. et al. Plasticity of Mitochondrial DNA Inheritance and its Impact on Nuclear Gene Transcription in Yeast Hybrids. Microorganisms 8, 494 (2020).

57. Szabó, A., Antunovics, Z., Karanyicz, E. & Sipiczki, M. Diversity and Postzygotic Evolution of the Mitochondrial Genome in Hybrids of Saccharomyces Species Isolated by Double Sterility Barrier. Front Microbiol 11, 838 (2020).

58. Hénault, M., Marsit, S., Charron, G. & Landry, C. R. Hybridization drives mitochondrial DNA degeneration and metabolic shift in a species with biparental mitochondrial inheritance. Genome Res 32, 2043–2056 (2022).

59. Chen, S., Zhou, Y., Chen, Y. & Gu, J. fastp: an ultra-fast all-in-one FASTQ preprocessor. Bioinformatics 34, i884–i890 (2018).

60. Li, H. & Durbin, R. Fast and accurate short read alignment with Burrows–Wheeler transform. Bioinformatics 25, 1754–1760 (2009).

61. Scannell, D. R. et al. The Awesome Power of Yeast Evolutionary Genetics: New Genome Sequences and Strain Resources for the Saccharomyces sensu stricto Genus. G3 Genes|Genomes|Genetics 1, 11–25 (2011).

62. Kent, W. J., Baertsch, R., Hinrichs, A., Miller, W. & Haussler, D. Evolution’s cauldron: Duplication, deletion, and rearrangement in the mouse and human genomes. Proceedings of the National Academy of Sciences 100, 11484–11489 (2003).

63. DoBlastzChainNet.pl - genomewiki. https://genomewiki.ucsc.edu/index.php/DoBlastzChainNet.pl.

64. Morrone, J. J. <strong>Biogeographical regionalisation of the Neotropical region</strong>. Zootaxa 3782, 1–110 (2014).

65. earthexplorer.usgs.gov. U.S. Geological Survey (1996).

66. Fick, S. E. & Hijmans, R. J. WorldClim 2: new 1□km spatial resolution climate surfaces for global land areas. Intl Journal of Climatology 37, 4302–4315 (2017).

67. García, E. Modificaciones al Sistema de Clasificación Climática de Köppen. (Instituto de Geografía, UNAM, México, 1973).

68. DePristo, M. A. et al. A framework for variation discovery and genotyping using next-generation DNA sequencing data. Nat Genet 43, 491–498 (2011).

69. Lachance, M.-A. Yeast communities in a natural tequila fermentation. Antonie van Leeuwenhoek 68, 151–160 (1995).

70. Ortiz, E. M. vcf2phylip v2.0: convert a VCF matrix into several matrix formats for phylogenetic analysis. zenodo DOI:10.5281/zenodo.2540861 (2019).

71. Stamatakis, A. RAxML version 8: a tool for phylogenetic analysis and post-analysis of large phylogenies. Bioinformatics 30, 1312–1313 (2014).

72. Alexander, D. H., Novembre, J. & Lange, K. Fast model-based estimation of ancestry in unrelated individuals. Genome Res. 19, 1655–1664 (2009).

73. McKenna, A. et al. The Genome Analysis Toolkit: A MapReduce framework for analyzing next-generation DNA sequencing data. Genome Res. 20, 1297–1303 (2010).

74. Danecek, P. et al. The variant call format and VCFtools. Bioinformatics 27, 2156–2158 (2011).

75. Cock, P. J. A. et al. Biopython: freely available Python tools for computational molecular biology and bioinformatics. Bioinformatics 25, 1422–1423 (2009).

76. Marr, R. A., et al. Whole genome sequencing of Canadian Saccharomyces cerevisiae strains isolated from spontaneous wine fermentations reveals a new Pacific West Coast Wine clade. G3 Genes|Genomes|Genetics 13, jkad130 (2023).

