## Supplementary Figures for "Widespread *Saccharomyces* hybridization in agave fermentations arises repeatedly from asymmetric parental diversity"

Supplementary Figure 1

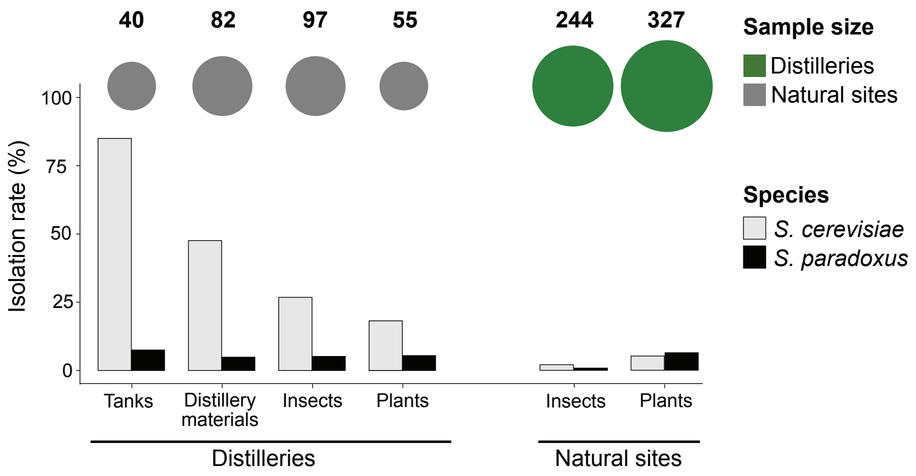

**Supplementary Figure 1**. Isolation of *Saccharomyces* spp. from the three geographical areas of Mexico where targeted sampling was performed. The bar plot shows the isolation rates as the percentage of samples that yielded *S. cerevisiae* (grey bars) or *S. paradoxus* (black bars) confidently identified with MALDI-TOF (score > 1.5) from each substrate type in distilleries and natural sites. Sample size of each category is shown on top.

### Supplementary Figure 2

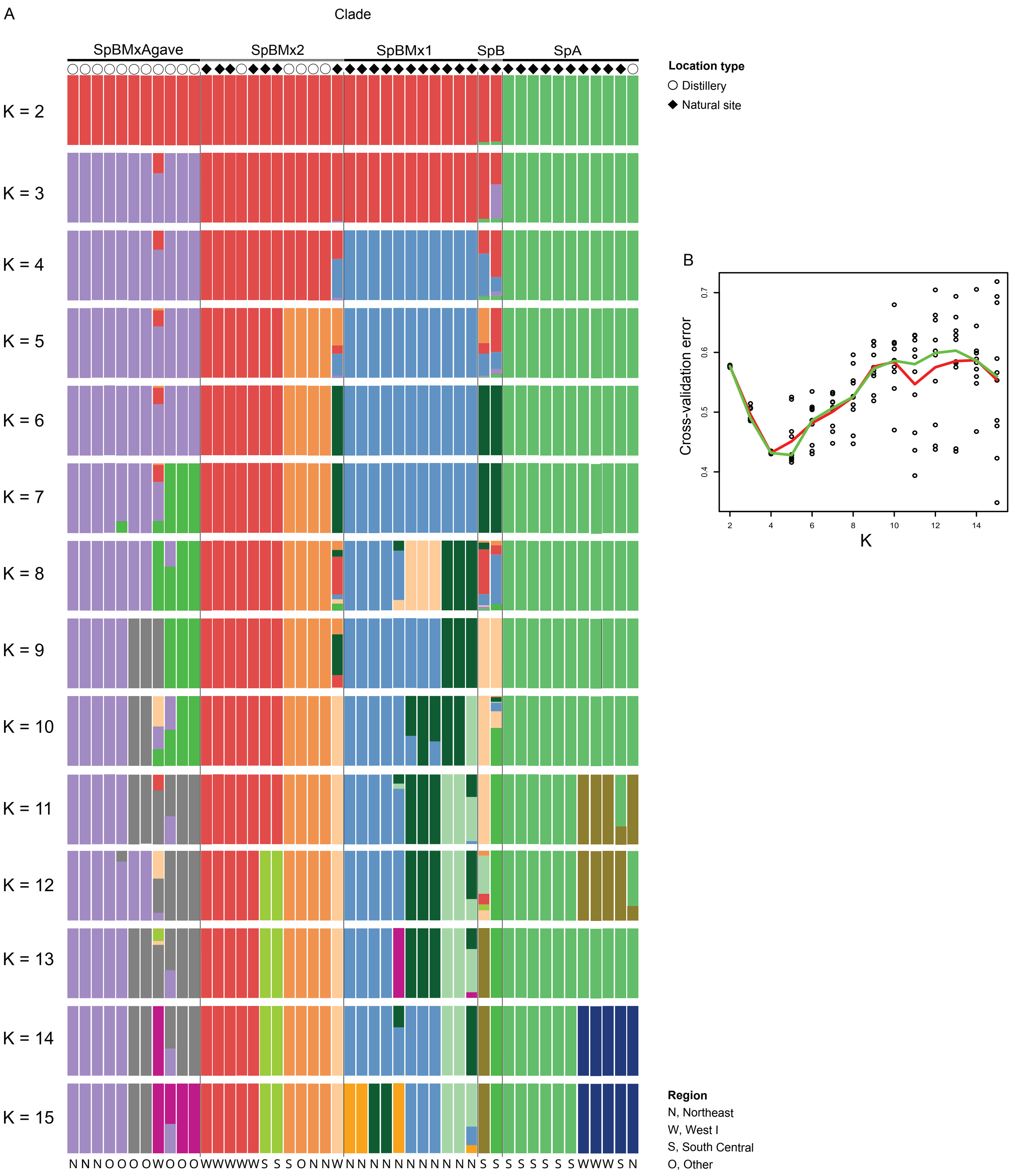

**Supplementary Figure 2**. Extended ADMIXTURE analysis of the *S. paradoxus* strains identified in the targeted sampling. A) ADMIXTURE plots for 47 *S. paradoxus* strains from $K = 2$ to $K = 15$, with 86,448 SNPs (previously LD pruned). B) ADMIXTURE cross-validation errors for $K = 2$ to $K = 15$, in a matrix of 86,448 SNPs (previously LD pruned) and 47 *S. paradoxus* strains. The red line represents the mean, whereas the green line represents the median.

Supplementary Figure 3

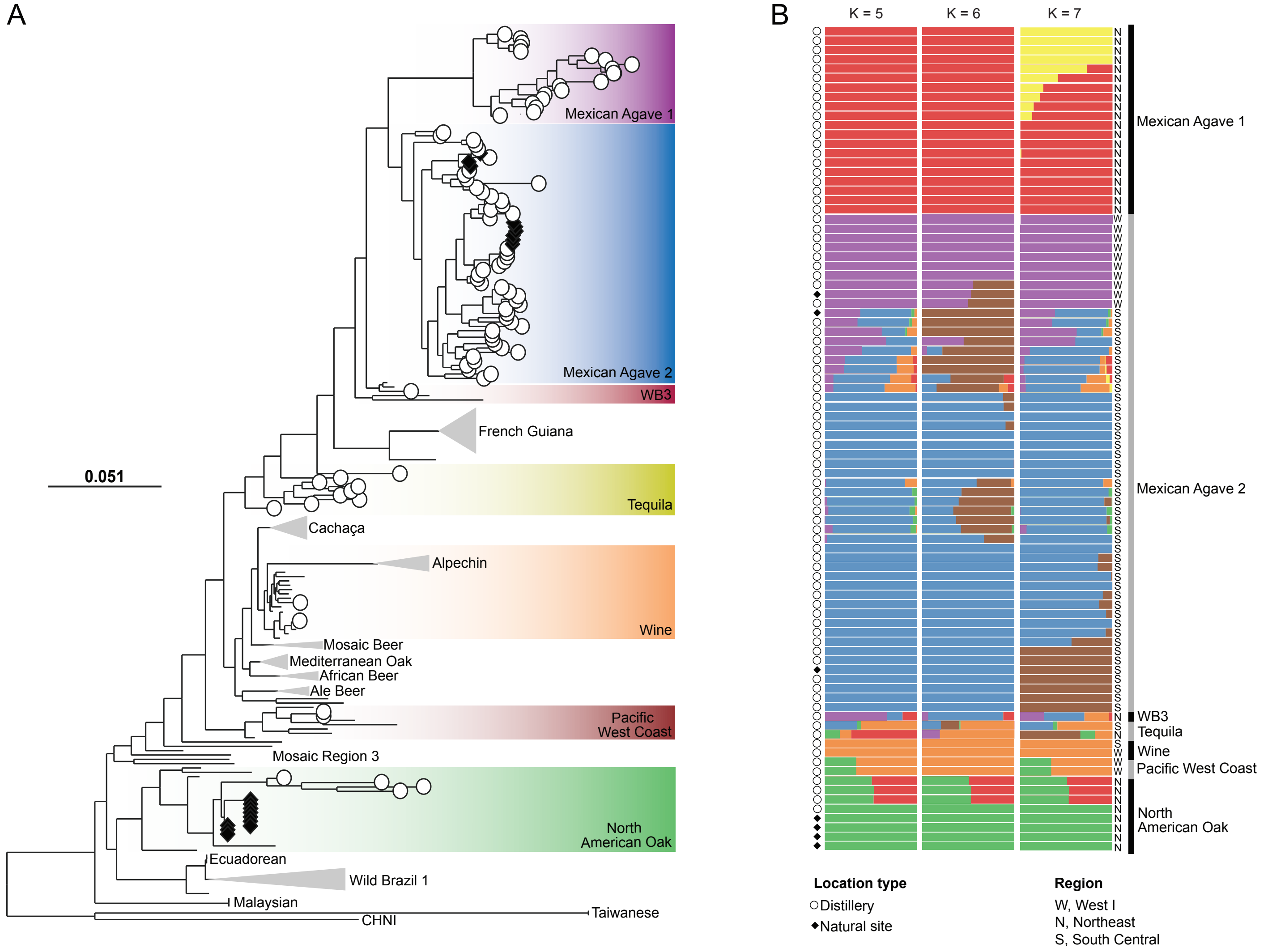

**Supplementary Figure 3**. Relationships of *S. cerevisiae* strains from agave spirit distilleries and natural sites in Mexico. **(A)** Maximum likelihood tree using 870,190 biallelic SNPs of 203 strains. Strains from agave spirit distilleries include 42 isolates from this study, 56 from [10], and 8 from [21]. All 19 *S. cerevisiae* isolates from natural sites in Mexico are from this study. The main clades of the world are indicated following the nomenclature of Peter and DeChiara, *et al.*[21] (with the exception of Wild Brazil 1 and 3 from[8] and Pacific West Coast from[76]); the Neotropical cluster was described by Avelar-Rivas *et al.*[10]. Only clades that include strains from this study or Avelar-Rivas *et al.*, 2026 are colored. Agave spirit producing regions only include the three regions visited during the targeted sampling in this study (Northeast, South Central and West I). Location type is indicated with white circles (○, distilleries) or black diamonds (♦, natural sites) at the tip of each branch**. (B)** Population assignment analysis using ADMIXTURE. Region is shown at the right; location type is indicated with white circles for distilleries and black diamonds for natural sites at the left. Assignation results for K = 2 to 15 are shown in **Supplementary Figure 4**.

### Supplementary Figure 4

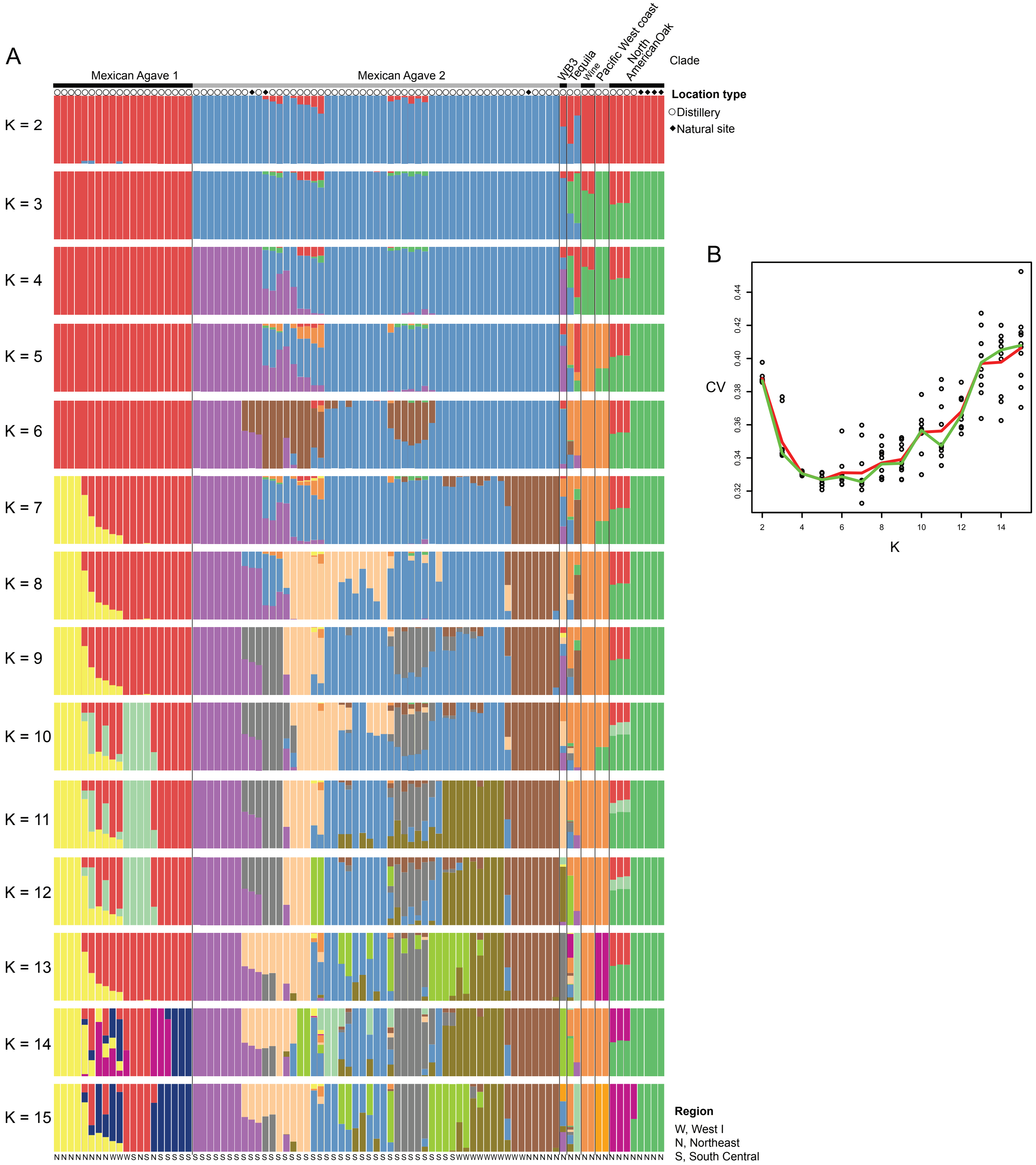

**Supplementary Figure 4**. ADMIXTURE analyses of *S. cerevisiae* strains from agave-spirits distilleries and natural sites of Mexico collected during the targeted sampling effort. **(A)** ADMIXTURE plots of 88 *S. cerevisiae* strains from K = 2 to K = 15. **(B)** ADMIXTURE cross-validation errors for K=2 to K=15, in a matrix of 108,636 SNPs (previously LD pruned) and 88 *S. cerevisiae* strains. The red line represents the mean, whereas the green line represents the median.

Supplementary Figure 5

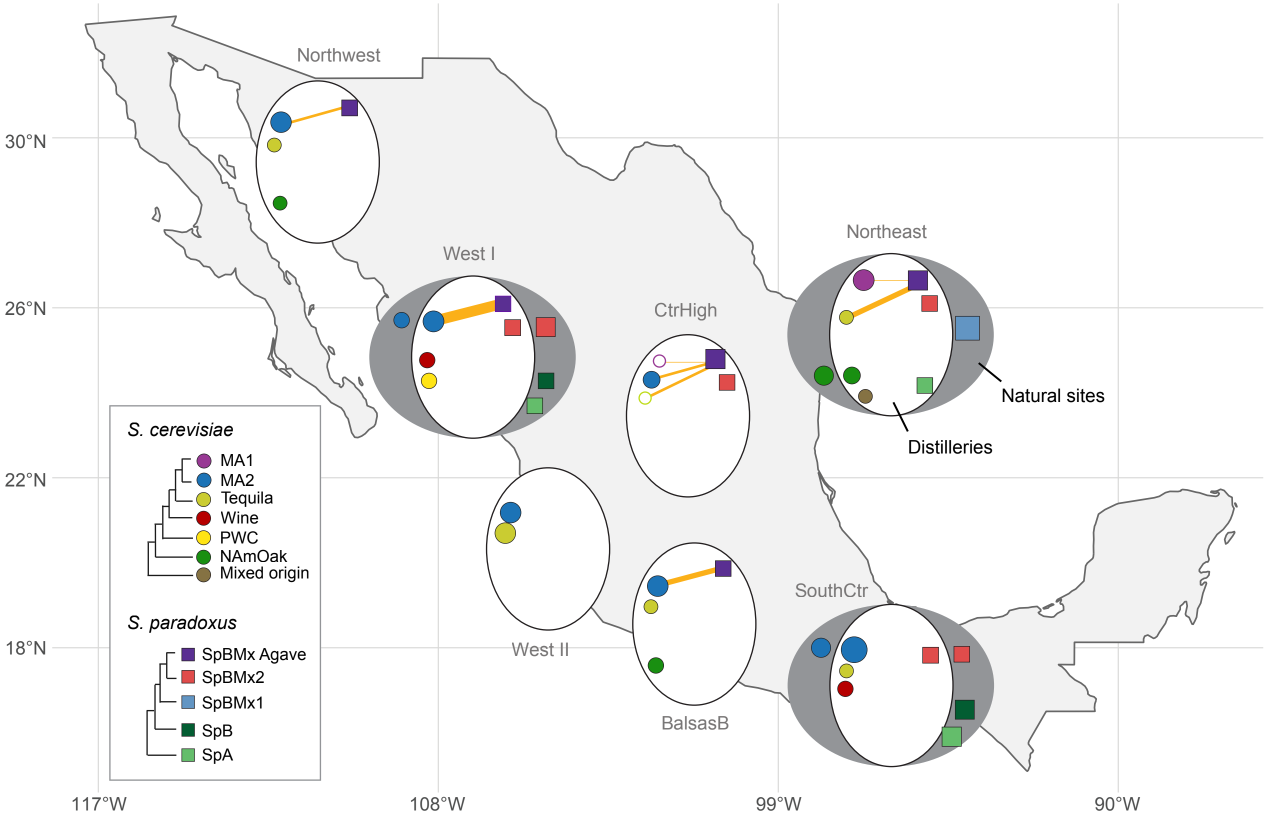

**Supplementary Figure 5**. Geographic distribution of *S. cerevisiae* and *S. paradoxus* populations associated with agave fermentations and natural sites in Mexico. Circular insets summarize the occurrence of *S. cerevisiae* (circles) and *S. paradoxus* (squares) subpopulations in each region sampled (Central Highlands [CtrHigh]; South Central [SouthCtr]; Balsas Basin [BalsasB]). Colors indicate the subpopulation assignments shown in the schematic trees at left. Symbols located inside the white circles represent strains isolated from distilleries, whereas symbols outside the circles correspond to strains recovered from natural sites that were sampled only in the West I, Northeast, and South Central regions. Symbol size is scaled according to the number of isolates belonging to each population within a region, with the visual scale capped at $n$=30; groups with more than 30 isolates are therefore shown at the maximum symbol size. Actual isolate counts range from 1 to 76. Red connecting solid lines indicate the occurrence of interspecific hybrids between *S. cerevisiae* and *S. paradoxus* subpopulations. Line thickness is scaled according to the number of independent hybrid strains inferred for each parental combination, with the visual scale capped at $n$=30; combinations with more than 30 hybrids are therefore shown at the maximum line width. Actual hybrid counts range from 1 to 32. Empty symbols connected by yellow lines indicate inferred parental populations for which no non-hybrid parental isolate was recovered from the corresponding region.

### Supplementary Figure 6

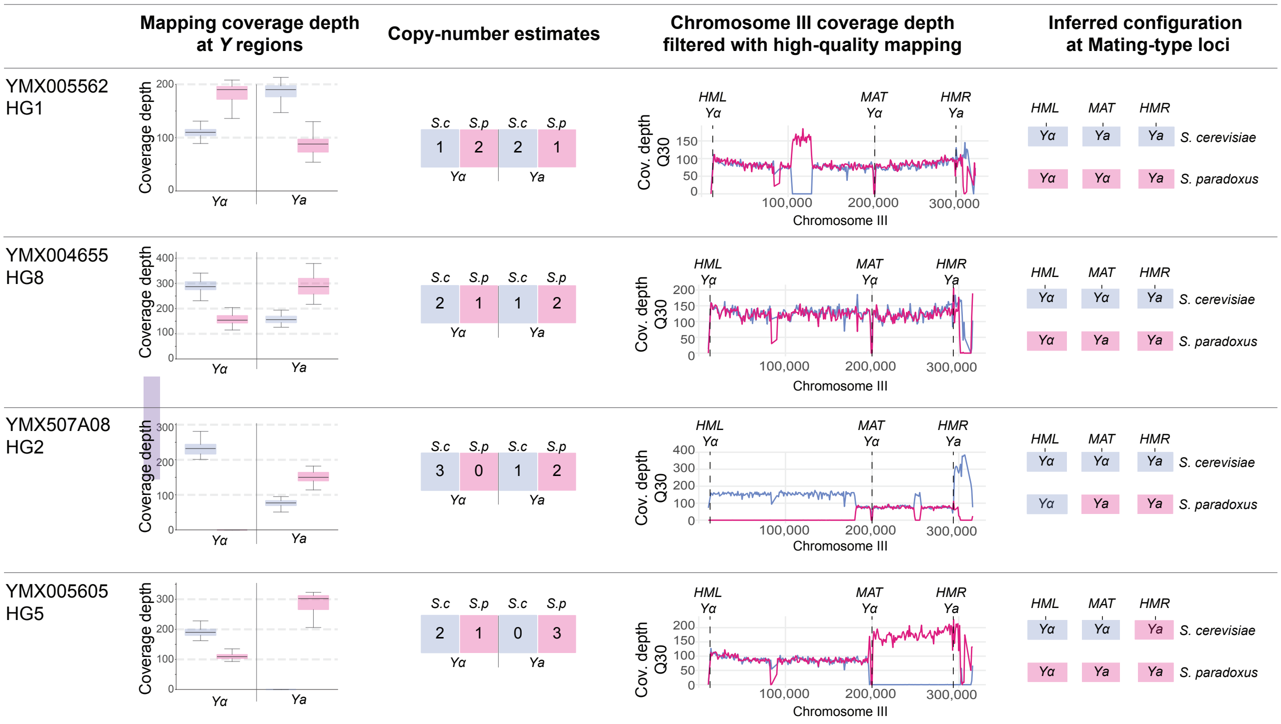

**Supplementary Figure 6.** Mating-type loci configurations in diploid Hybrid Groups support independent hybridization events. The first column shows the per-base coverage across the mating-type-specific *Ya* and *Yα* regions of both parental species. No mapping-quality filter was applied for this analysis allowing multimapping reads with equally good alignments to be randomly assigned to one of the equally-scoring positions. The coverage of the two identical *Yα* regions was added. Boxplots show the median and interquartile range, with whiskers extending 1.5x the ICR. In column two**,** copy-number estimates show relative copy numbers of *Ya* and *Yα* regions in each species. This was calculated by dividing each median by the smallest non-zero median coverage within each isolate and rounding to the nearest integer. The third column shows the mean 1kb windowed coverage after filtering as described in Methods across Chromosome III. The *HML*, *MAT* and *HMR* loci are shown, highlighting the location of the *Ya* or *Yα* in both subgenomes in the concatenated reference. Coverage at the mating-type cassettes themselves is depleted at Q30 filtering because reads from these homologous regions frequently map ambiguously; therefore, local ancestry was inferred from the pattern of allelic balance at the flanking regions. The last column shows the inferred mating-type locus configuration, representing the most parsimonious arrangement consistent with the estimated *Y*-region copy numbers and the chromosome III ancestry together with LOH patterns shown in column three.

### Supplementary Figure 7

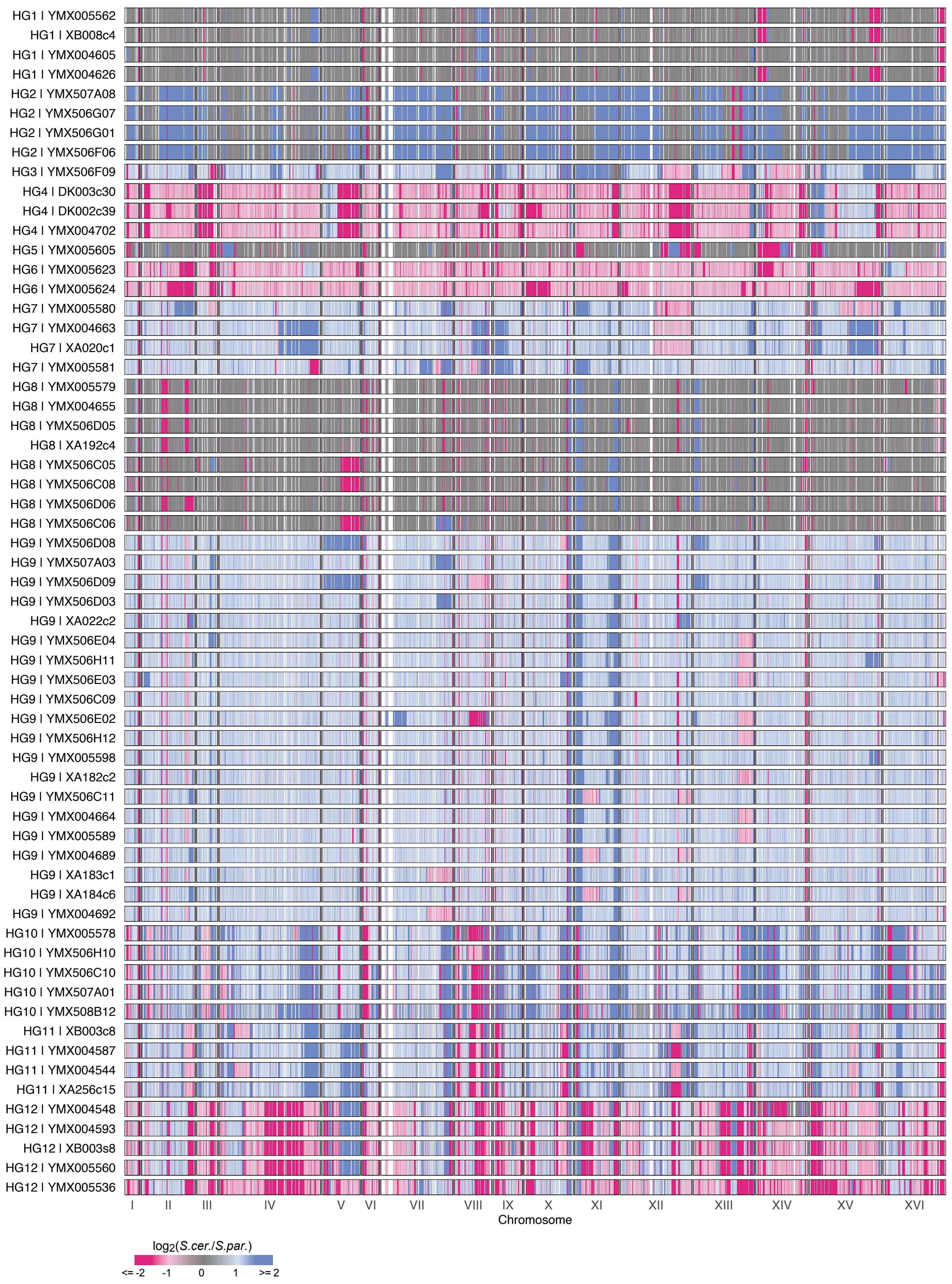

**Supplementary Figure 7.** Relative subgenome coverage of the 61 hybrids sequenced in this study. The $\log_2$ ratio of *S. cerevisiae* to *S. paradoxus* sequencing depth, calculated in 1-kb windows with at least 80% of bases aligned between the parental reference genomes is used as a proxy for subgenome balance. The resulting profiles reveal distinct patterns among hybrid groups (HGs), as well as variation within individual groups. Hybrids are ordered as in Fig. 4B, with members of each HG displayed consecutively.
